# Polyphosphate as a Novel Regulator of Super-Enhancer Complexes: Disruption of Phase Separation and Gene Expression

**DOI:** 10.1101/2024.12.12.628164

**Authors:** Zhiyun Yang, Zongchao Jia

## Abstract

Inorganic polyphosphate (polyP) is a highly conserved linear polymer of orthophosphate, present across nearly all living organisms. Its functional diversity, particularly through recently discovered post-translational modifications, has garnered increasing scientific interest. However, its role in gene regulation, especially in mammalian systems, remains unexplored. In this study, we present findings that demonstrate that polyP multi-targets the super-enhancer complex to regulate gene expression, marking a novel mechanism of transcriptional control. We show that polyP modifies key components of the complex, including the Mediator subunit MED1, the coactivator Bromodomain-containing protein 4 (BRD4), and the transcriptional regulator Yin Yang 1 (YY1). This polyP-mediated modification not only disrupts their ability to undergo phase separation but also downregulates the expression of MED1 and BRD4, as well as impairs the nuclear localization of YY1. Furthermore, polyP inhibits YY1 dimer formation and disrupts YY1-mediated DNA looping, leading to suppressed gene expression. These discoveries highlight polyP as a novel regulator of super-enhancer-driven gene transcription and provide new insights into the broader role of polyP in epigenetic regulation. This work uncovers a previously unknown layer of gene control, positioning polyP as a critical player in transcriptional regulation through its interaction with transcription factors and coactivators.

**Graphical abstract:** 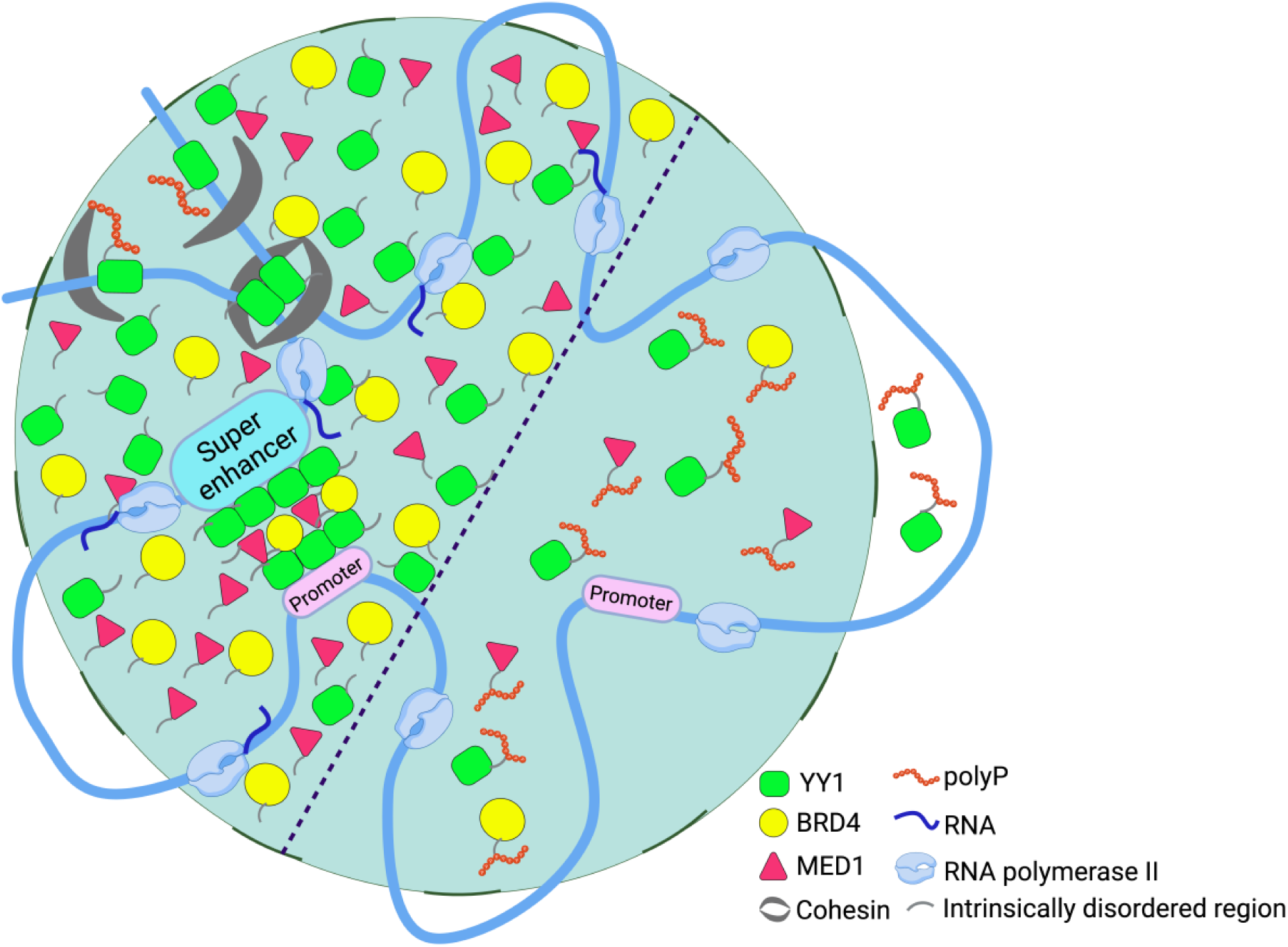

**Highlights:**

- PolyP modifies key components of the super-enhancer complex BRD4, MED1 and YY1 by targeting their His-rich, Lys-rich or PASK sequences
- PolyP disrupts phase separation of BRD4, MED1 and YY1
- PolyP impairs YY1’s recruitment of BRD4 and MED1 and disrupts nuclear localization, dimerization, DNA looping and DNA binding to YY1
- PolyP attenuates gene expression mediated by YY1 and BRD4

## Introduction

PolyP is a linear polymer connected by energy-rich phosphoanhydride bonds, ranging from three to a few hundreds of repeating inorganic phosphate units. In bacteria, polyP chains (100-1000 residues) are synthesized by polyphosphate kinase (PPK) enzymes, PPK1 and PPK2;^1–3^ and degraded by exopolyphosphatase (PPX).^4^ PolyP and PPKs from bacteria such as *Pseudomonas aeruginosa* are associated with bacterial virulence such as biofilm formation and motility.^5–7^ In yeast, vacuolar transporter chaperone complexes (VTC) are found to be responsible for polyP synthesis;^8,9^ while in higher eukaryotes, lacking PPK homologues, the source of polyP (50-150 residues) remains mysterious.^10,11^ However, polyP in human platelets can accumulate up to 130 mM in dense granules,^12^ playing an important role in inflammation and blood clotting.^13^ Several mammalian enzymes have been identified as polyphosphatase candidates including H-Prune,^14^ Nudix family members DIPP1 (also call Nudt3),^15,16^ DIPP2 and DIPP3.^15^

A remarkable function of polyP in eukaryotes is its post-translational modification on proteins. The first modification was discovered on yeast nuclear signal recognition 1 (Nsr1) and topoisomerase 1 (Top1), where polyP attached to lysine residues within a PASK (defined as a stretch of 20 amino acids with at least 75% D/E/S and at least one K).^17,18^ This impaired Top1 enzymatic activity and interfered its interaction with Nsr1.^17^ Downey lab took advantages of this domain to identify 29 polyP targets, 23 from yeast^18,19^ and 6 human proteins.^18^ Among which, 15 yeast targets formed a conserved network involved in ribosome biogenesis where they found that polyP is playing a role in regulating ribosome function.^18^ Subsequent study using microchip identified another 4 polyP targets from human cells without containing obvious PASK domain,^20^ which promotes further studies to define polyP targeting domain. Recently, we showed that proteins containing consecutive histidine residues are targeted by polyP and documented at least 30 human and yeast histidine repeat proteins bound by polyP.^21^ Importantly, the discovery of the binding between polyP and His-repeat protein has expanded polyP functions including disrupting phase separation and phosphorylation activity of the human protein kinase DYRK1A and inhibiting the activity of the transcription factor MafB,^21^ indicating polyP modification could work as a potential regulatory mechanism. Given the PTM-like characteristics, it has been termed histidine polyphosphate modification or HPM. Several groups have studied the impact of polyP in mammalian cells by ectopically expressing PPX or PPK. Wang et al.^22^ first showed that polyP activates the mammalian target of rapamycin (mTOR) through expressing yeast PPX1 in MCF-7 mammary cancer cells. This is further confirmed by Hassanian et al.^23^ revealing that polyP activates mTORC1 and mTORC2 to exert its proinflammatory effect in endothelial cells by adding polyP to the culture media. Bondy-Chorney et al.^24^ showed a broad change elicited by polyP through overexpressing *E. coli* PPK in human cells including transcriptome and proteome reprogramming, activation of ERK1/2-EGR1 signaling pathway and relocalization of nuclear/cytoskeleton proteins DEK, TAF10, GTF2I and eIF5b. Other roles of polyP were also illustrated in mitochondria metabolism^25^ and assisting DNA damage repair.^26^ More recently, we reported that, in addition to the PASK motif, lysine-rich sequences in proteins also undergo polyP modification (KPM), akin to histidine polyphosphate modification (HPM).^27^

The Young lab was the first to propose the concept of super-enhancers, describing them as complexes consisting of clusters of enhancers that are cooperatively occupied by a high density of master transcriptional regulators and Mediator.^28^ Comparing to typical enhancers, super-enhancers can harbor about 10-fold higher density of these factors, which drive robust expression of genes with prominent roles in human cell identity.^28–30^ Super-enhancers are enriched at oncogenes, driving their high-level expression and contributing to tumorigenesis. As a result, cancer cells are addicted to super-enhancer driven transcriptional programs to maintain their identity.^29,31–33^ This dependence suggests that super-enhancers could serve as valuable biomarkers for cancer diagnosis and therapeutic targeting. The presence of transcriptional coactivators such as BRD4 and MED1, which are enriched at super-enhancers, has been widely used to define and characterize these regulatory elements.^28,29,34,35^ These two coactivators can form phase-separated condensates to compartmentalize and concentrate transcription apparatus, which are involved in gene activation.^35,36^ Lovén et al.^30^ have shown that inhibition of BRD4 with BET-bromodomain inhibitor JQ1 caused preferentially loss of transcription at super-enhancer associated genes. Cho et al.^37^ revealed that Mediator is sensitive to transcriptional inhibitors which indicates that Mediator is dynamically interact with transcription factors to initiate transcription.

Transcription factors play an essential role in cell growth, development and differentiation through activation, repression and modification of gene expression.^38,39^ Yin Yang 1 (YY1) is one such multifunctional ubiquitous transcription factor. It is expressed throughout mammalian cells and modulates many crucial biological and cellular processes. Nuclear localization is essential for YY1’s transcriptional regulation and control,^40^ where it binds to its consensus motif (GCCGCCATTTTG) to regulate gene expression.^41^ Histidine cluster of YY1 is responsible for its nuclear localization and formation of liquid-liquid phase separation, which is important for compartmentalizing coactivators and enhancer elements like BRD4 band MED1.^42,43^ Dimerization of YY1 is essential for its gene regulation. YY1 was reported to form homodimers and bind to consensus sequence present in enhancers and promoters to mediate physical loop formation between enhancers and promoters in super-enhancer.^44^ In addition, YY1 dimers was also reported to bind DNA guanine quadruplexes (G4) structure, stable secondary structures of DNA that plays important role in DNA replication and transcription,^45^ to mediate long-range DNA looping and regulate gene expression.^46^

Here, we identified polyP as a putative regulator of super-enhancer through its multi-targeting effects on key proteins within the complex, MED1, BRD4 and YY1. Our findings demonstrate that polyP modification of these proteins not only destroys their phase separation but also decreases the expression of MED1 and BRD4, while blocking nuclear localization of YY1. Furthermore, polyP interferes with critical functions of YY1, including its binding with YY1 consensus sequence and G4 structure, dimerization, and YY1-mediated DNA looping. These disruptions collectively inhibit super-enhancer-driven gene transcription.

## Results

### Polyphosphate modifies super-enhancer complex markers, BRD4 and MED1

Twelve polyP targets implicated in ribosome biogenesis have been identified and are localized to the nucleolus, one of the three major chromatin subcompartments.^19,47^ This suggests that polyP might work as a regulator of gene expression by targeting chromatin subcompartments. Surprising, we found 4 out of 10 chromatin subcompartment marker proteins which can undergo liquid-liquid phase separation^47^ contain PASK domain: NPM1, BRD4, MED1 and HP1. Thus, we assessed whether these four proteins could be modified by polyP using NuPAGE gel. The results confirmed that polyP can modify BRD4 (Fig. 1A) and MED1 (Fig. 1B), as evidenced by an electrophoretic mobility shift in the presence of polyP_700_. However, although there are two PASK domains within NPM1 and HP1 (Fig. S1A and S1B), polyP was unable to modify them (Fig. S1C). Since the enzyme(s) responsible for polyP synthesis is unknown, PPK expression in mammalian cells has been widely used to study the functions of polyP.^21,24^ To test the effects of polyP on BRD4 and MED1, we transiently transfected either an empty vector (mCherry) or a vector expressing the *Pseudomonas aeruginosa ppk1* gene under the cytomegalovirus (CMV) promoter in HeLa cells. The PPK1 expression resulted in accumulation of polyP in HeLa cells (Fig. 1C). To investigate the impact of polyP accumulation on super-enhancer complex, we examined protein changes of super-enhancer markers by western blot. We observed a significant decrease both in BRD4 and MED1 proteins as compared to normalized control (Fig. 1D and 1E).

**Figure 1.**
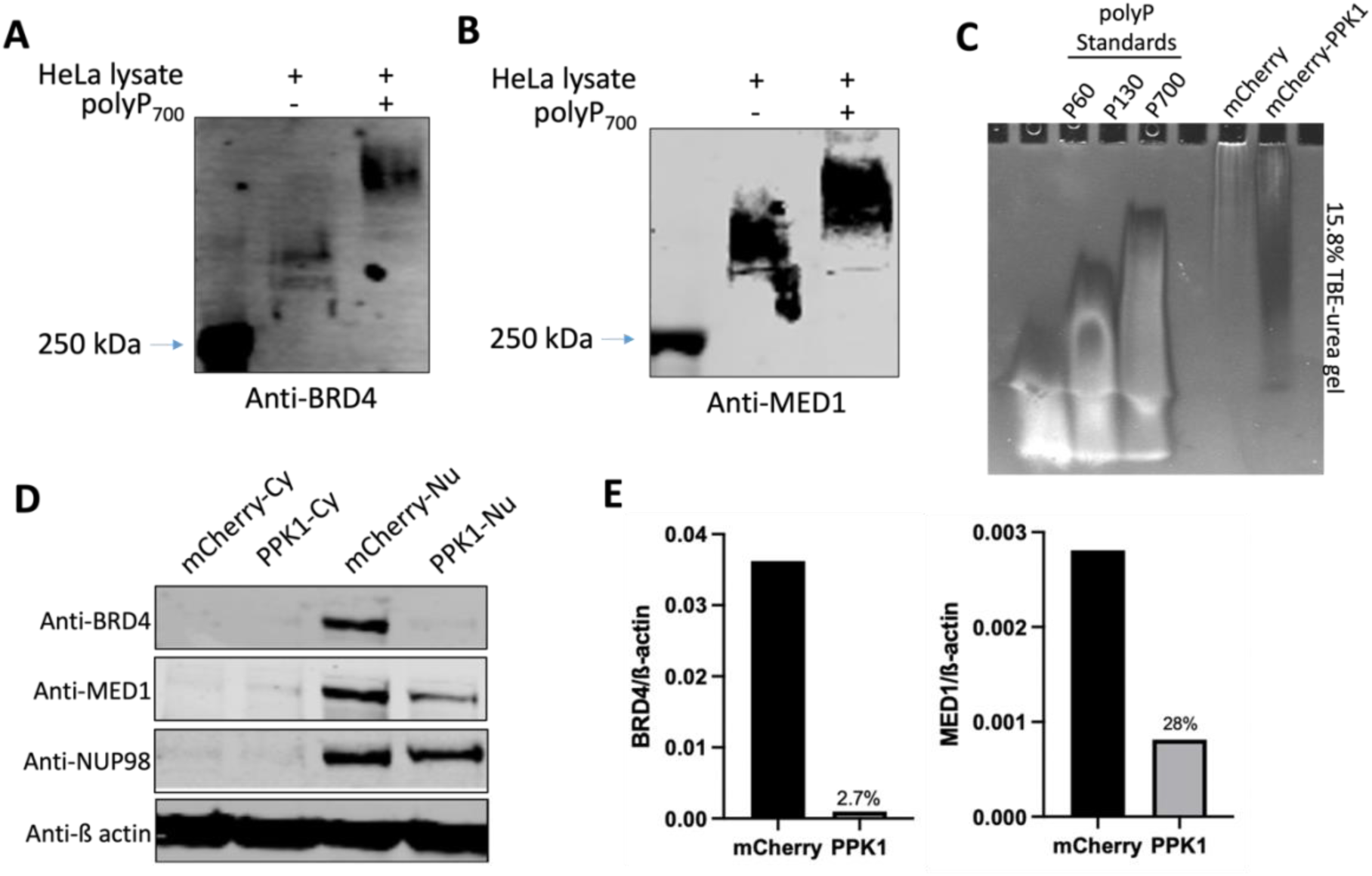
polyphosphate modifies MED1 and BRD4. (A and B) PolyP modification of BRD4 (A) and MED1 (B). HeLa cells lysate with/without the addition of polyP_700_ were analyzed via NuPAGE followed by western blot with antibodies against BRD4 (A) and MED1 (B). Images are representative of n = 3. PolyP_700_ was present at 5 mM unless otherwise indicated. (C) PolyP overproduction in HeLa cells. polyP extractions from HeLa cells transfected with mCherry or mCherry-PPK1 were analyzed on a 15.8% Tris-borate-EDTA (TBE)-urea gel stained with DAPI. PolyP standards (60, 130 and 700 phosphates) are presented for comparison (n = 3). (D) Western blot analysis after SDS-PAGE of fractions from mCherry or mCherry-PPK1 expressing HeLa cells with antibodies against proteins localized to specific fractions: cytoplasm (Cy) and nuclear (Nu). Western blot shown is a representative image of n = 3. (E) Quantification of changes in nuclear protein levels of BRD4 and MED1 relative β-actin from (D).

### Polyphosphate disrupts liquid-liquid phase separation of BRD4-PIR and MED1-PIR

BRD4 and MED1 contain large intrinsically disordered region (IDR), which has been shown to enable them to undergo liquid–liquid phase separation at super-enhancers.^35^ BRD4 has both PASK domain and 6 histidines which are polyP targeting sequences (Fig. 2A and 2B); and MED1 has PASK domain as well (Fig. 2A and 2B). Importantly, the PASK and histidine-rich regions, along with their neighboring sequences in BRD4 and MED1, are highly conserved across different vertebrate species (Fig. 2B). Therefore, we designate these conserved sequences as polyP interaction region (PIR): BRD4-PIR (amino acid 648-752) and MED1-PIR (amino acid 1458-1581) and selected them for further study. Using purified BRD4-PIR and MED1-PIR, we confirmed that polyP_700_ modifies BRD4 and MED1 through their PIR (Fig. 2C, S2A and S2B). Next, we examined the salt sensitivity of polyP modification on BRD4-PIR and MED1-PIR. We found high concentration of NaCl completely abolished the shift of BRD4-PIR and MED1-PIR at 200 mM and 400 mM respectively in NuPAGE gel (Fig. 2D). The IDRs of BRD4 and MED1 proteins are known to be involved in formation of phase-separated droplets *in vitro*.^35^ Therefore, we investigated whether the PIRs of BRD4 or MED1 can form such droplets *in vitro*. Purified recombinant GFP-fusion proteins (BRD4-PIR and MED1-PIR) were added to buffers containing 10% PEG-8000 (polyethylene glycol, molecular weight 8000), turning the solution opaque with the presence of salt concentration from 50 to 500 mM (Fig. S2C). While polyP disrupted these droplets formation in the presence of salt concentration up to 125 mM and 200 mM for BRD4-PASK and MED1-PASK respectively (Fig. 2E), higher concentration of salt did not have the same effect (Fig. 2D and 2E). These results indicate that the phase separation of BRD4 and MED1, previously shown to occur at super-enhancers, could be mediated by polyP. Additionally, this process may be fine-tuned by ionic strength.

**Figure 2.**
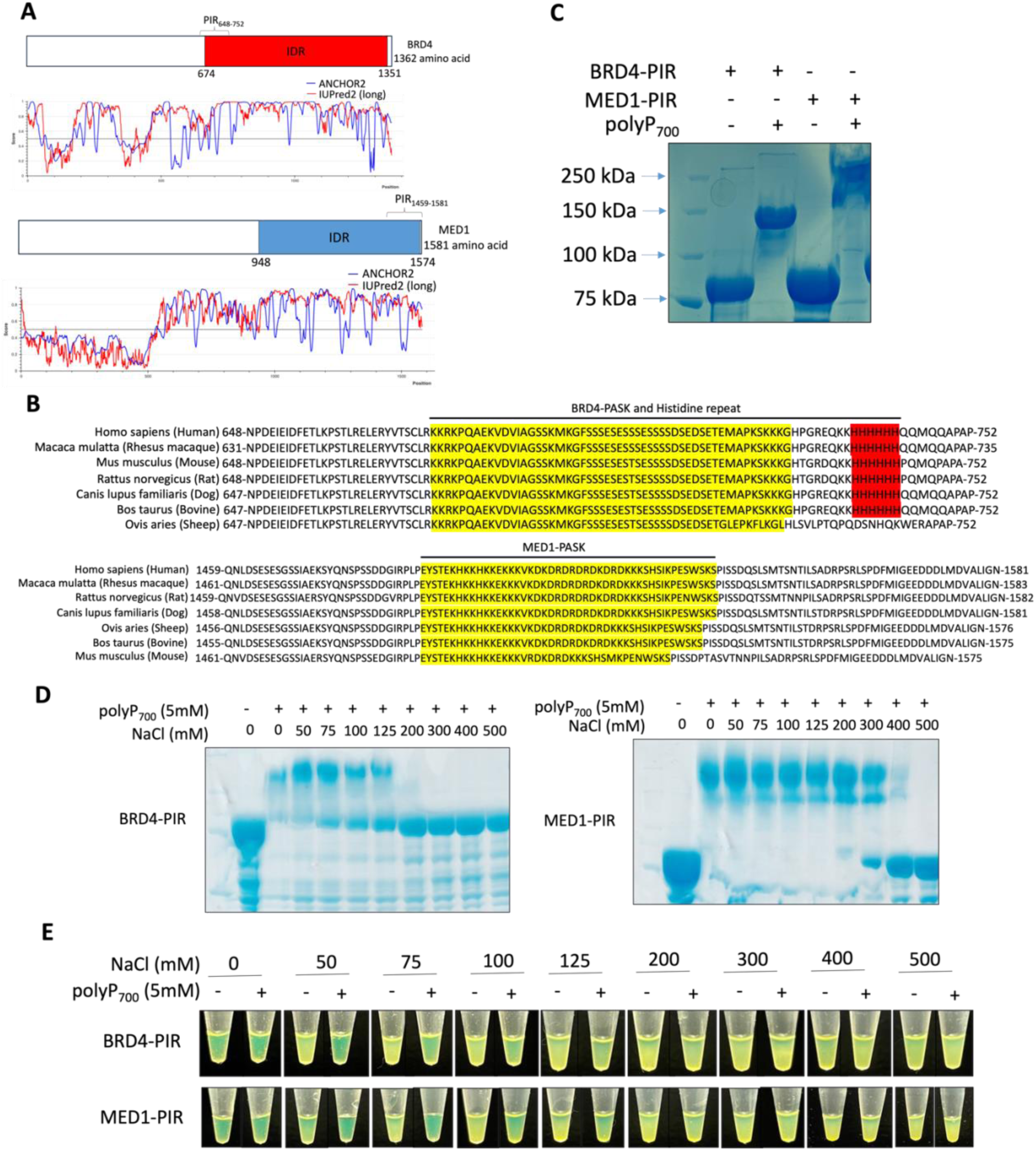
Polyphosphate disrupts liquid-liquid phase separation of BRD4 and MED1 in a salt-dependent manner *in vitro*. (A) Domain graphs of BRD4 and MED1 PIRs; IDRs prediction based on ANCHOR2 and IUPred2 algorithms. Scores >0.5 indicate disorder. (B) PIRs of BRD4 and MED1 proteins from different vertebrates. (C) Coomassie-stained NuPAGE analysis showing polyP-mediated shift of purified MBP-GFP tagged BRD4-PIR and MED1-PIR. (D) Coomassie-stained NuPAGE analysis of polyP_700_ modification of purified MBP-GFP tagged BRD4-PIR (Left) and MED1-PIR (Right) with the indicated concentration of salt (n=3). (E) Phase separation of purified MBP-GFP tagged BRD4-PIR and MED1-PIR with and without polyP_700_ under the indicated concentration of salt. Tubes containing MBP-GFP tagged BRD4-PIR or MED1-PIR in the buffer containing PEG8000 (n=3).

### Polyphosphate impairs nuclear localization and liquid-liquid phase separation of YY1

We next investigated whether polyP could serve as a regulator of super-enhancer-mediated gene regulation. YY1 is one of the transcription factors governed by super-enhancer regulatory networks. Our lab found that YY1 undergoes HPM, evidence by polyP-mediated shift on NuPAGE gel (Fig. S3A and S3B).^21^ A stretch of 11 histidines (aa 70–80) within the acidic transactivation domain (TAD) (Fig. 3A) is thought to be targeted by polyP (Fig. S3C).^21^ To confirm this, we performed the polyP shift experiment and showed that purified TAD underwent HPM upon addition of polyP_700_ (Fig. 3B). Histidine-repeat is responsible for stimulating YY1 accumulation in nuclear speckles.^42^ Therefore, we further investigated the impact of polyP on YY1 activities. We first examined changes in YY1 protein levels in HeLa cells using western blot analysis. In the PPK1-expressing cells, the total YY1 levels were not affected (Fig. 3C) but the localization of YY1 to nuclear was impaired (Fig. 3D and 3E). Taking advantage of the partial localization of YY1 to the nucleus in the presence of polyP, we next tested if the polyP produced by PPK1 could enter the nucleus using NuPAGE gel. Our result showed that the polyP can enter nucleus, with the length of polyP appearing to shrink. This is indicated by the faster mobility of nuclear YY1 on NuPAGE compared to the cytoplasmic YY1 (Fig. 3F). We also investigated the impact of polyP on YY1 nuclear sparkle formation by co-transfecting vectors containing YY1 and PPK1 in HeLa cells. Fluorescence imaging results showed that polyP prevents nuclear sparkle formation of YY1 in PPK1-expressing HeLa cells (Fig. 3G). In addition, the imaging results also showed that polyP inhibits the nuclear localization of YY1 (Fig. 3G), which is consistent with the western blot results (Fig. 3D, 3E and 3F). Importantly, we found that polyP can destroy the pre-formed nuclear sparkle of YY1 when we treated the fixed HeLa cells expressing GFP-YY1 with polyP_700_ (Fig. 3H). Lastly, we tested if YY1 HPM is also salt dependent using NuPAGE gel. The results showed that the polyP_700_ can induce a shift in YY1 in the presence of salt concentrations of 0-300 mM (Fig. 3I). YY1 is known to form liquid-liquid phase separation *in vitro*.^43^ We next tested whether polyP_700_ can compromise *in vitro* phase separation of YY1. The results demonstrated that, in the presence of salt concentration between 50-200 mM, polyP_700_ effectively disrupted YY1 phase separation (Fig. 3J). These results collectively indicate that polyP may affect YY1’s transactivation functions.

**Figure 3.**
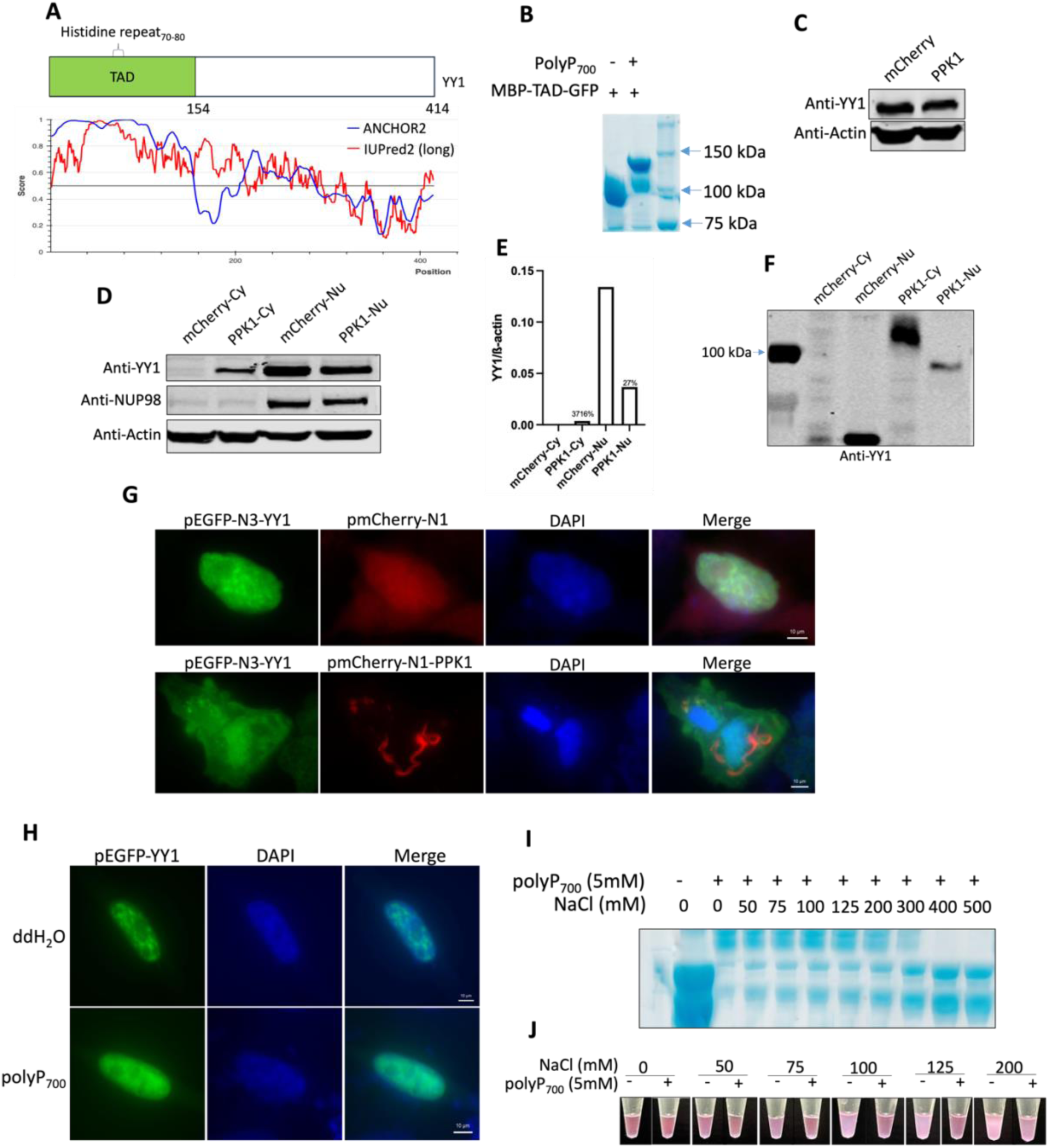
Polyphosphate abolishes the nuclear localization and phase separation of YY1. (A) Domain graphs of YY1 TAD; IDRs prediction based on ANCHOR2 and IUPred2 algorithms. Scores >0.5 indicate disorder. (B) Coomassie-stained NuPAGE analysis of purified MBP-GFP tagged YY1 TAD modified by polyP_700_. (C) Western blot analysis after SDS-PAGE using the YY1 antibody with β-actin as a loading control (n = 3). (D) Western blot analysis after SDS-PAGE of fractions from mCherry or mCherry-PPK1 expressing HeLa cells with antibodies against proteins localized to specific fractions: cytoplasm (Cy) and nuclear (Nu) (n=3). (E) Quantification of changes in cytoplasmic and nuclear protein levels of YY1 relative to β-actin from (D). (F) Western blot analysis after NuPAGE using the YY1 antibody (n=3). (G) PolyP overproduction in HeLa cells results in the loss of YY1 droplets. HeLa cells were co-transfected with GFP-YY1 and either mCherry or mCherry-PPK1 (n=3). (H) PolyP addition to the fixed GFP-YY1 expressing HeLa cells results in the loss of pre-formed YY1 droplets (n=3). (I) Coomassie-stained NuPAGE analysis of polyP_700_ modification of purified MBP-mCherry tagged YY1 with the indicated concentration of salt (n=3). (J) phase separation of purified MBP-mCherry tagged YY1 with and without polyP_700_ under the indicated concentration of salt. Tubes containing MBP-mCherry tagged YY1 in the buffer containing PEG8000 (n=3).

### Polyphosphate affects the co-localization of YY1 and BRD4 or MED1

To establish super-enhancer, transcription factors compartmentalize transcriptional coactivators, such as BRD4 and MED1, to mediate gene transcription.^35,36^ Previous study has shown that YY1 can colocalize with BRD4-IDR and MED1-IDR *in vitro*,^43^ which is potentially important for YY1-mediated gene transcription. Here, we investigated whether BRD4-PIR and MED1-PIR can colocalize with YY1 *in vitro*. Purified mCherry-YY1 was added to buffers containing 10% PEG-8000 with either purified GFP-BRD4-PIR or GFP-MED1-PIR. Our results showed that mCherry-YY1 forms droplets that overlap with droplets formed by GFP-BRD4-PIR (Fig. S2D) or GFP-MED1-PIR (Fig. S2E). Next, we assessed whether polyP prevents the colocalization of YY1 and BRD4-PIR or MED1-PIR. The results revealed that the droplets formed by YY1, BRD4-PIR and MED1-PIR are much smaller in the presence of polyP_700_ (Fig. 4A and 4B), while mCherry-YY1 fails to colocalize with GFP-BRD4-PASK (Fig. 4A) or GFP-MED1-PASK (Fig. 4B). These results support the idea that polyP may interfere with the recruitment of coactivators BRD4 and MED1 by YY1, thereby disrupting the formation of super-enhancers.

**Figure 4.**
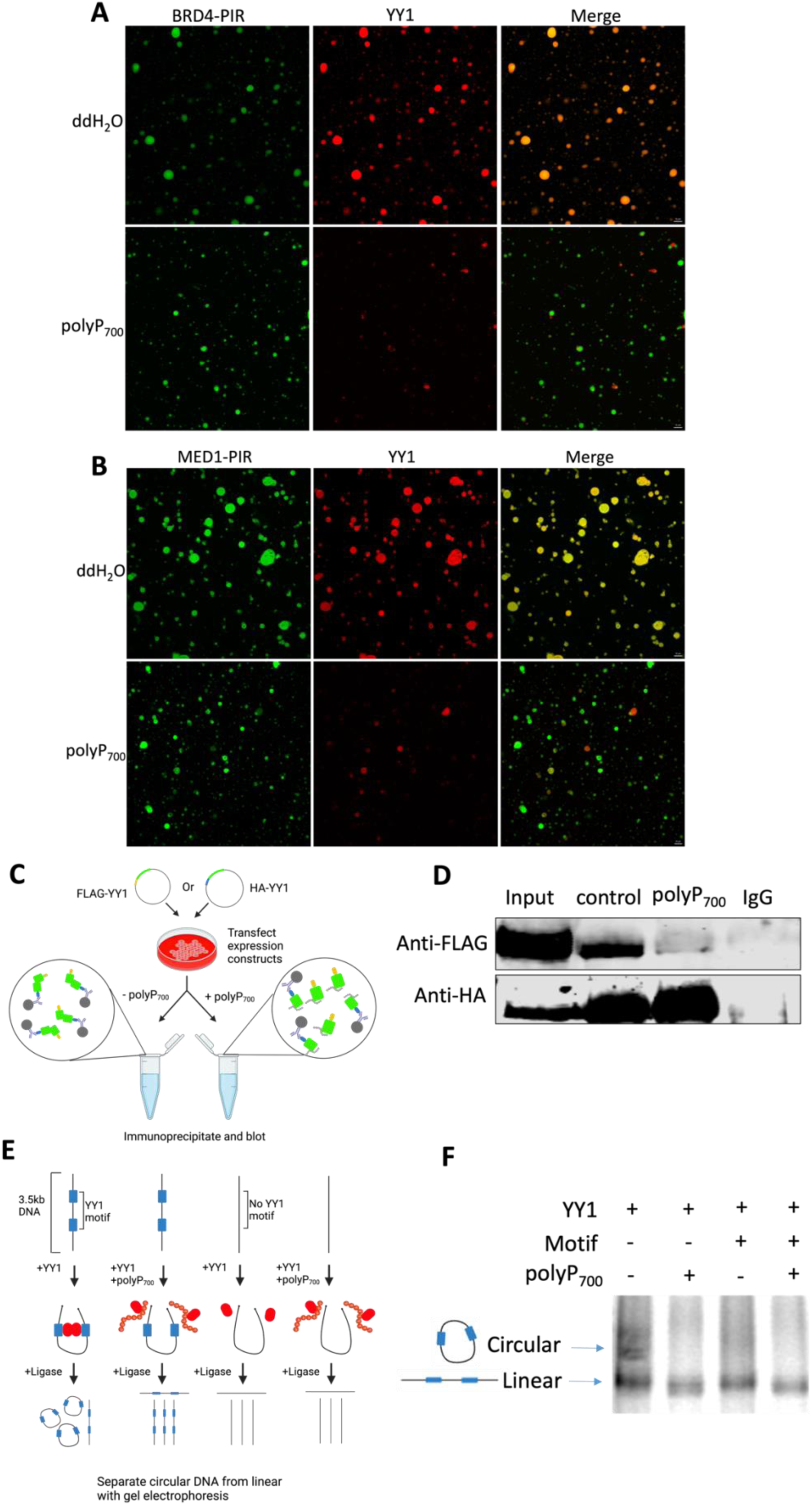
Polyphosphate inhibits YY1’s recruitment of BRD4-PIR and MED1-PIR, and impairs YY1 dimerization and DNA looping function. (A and B) PolyP_700_ affects the droplet formation of BRD4-PIR (A), MED1-PIR (B) and YY1 (A and B); and the ability of YY1 droplets to incorporate BRD4-PIR (A) or MED-PIR (B) proteins *in vitro*. MBP-GFP or MBP-mCherry fusion proteins were mixed in buffer containing PEG8000 and 100 mM NaCl. Indicated fluorescence channels are presented for each mixture (n=3). (C) Model depicting co-immunoprecipitation assay to detect YY1 dimerization with and without the addition of polyP_700_ in the resin mixture. (D) Western blot analysis after SDS-PAGE showing the ability of polyP_700_ to disrupt co-immunoprecipitation of FLAG-tagged YY1 and HA-tagged YY1 proteins from nuclear lysates prepared from transfected cells using antibodies against FLAG or HA (n=3). (E) Model depicting the *in vitro* DNA circularization assays used to detect YY1-mediated DNA looping interactions with/without the addition of polyP_700_. (F) Results of the *in vitro* DNA circularization assay visualized by gel electrophoresis showing the ability of polyP_700_ to disrupt YY1-mediated DNA loop formation.

YY1 acts as a dimer to bind to consensus sequence present in enhancers and promoters across cell types,^32,44^ which is essential for super-enhancer formation. To further investigate the mechanism by which polyP affects the recruitment of coactivators by YY1, we next assessed whether polyP impairs YY1’s function as chromatin regulator. Specifically, we investigated if polyP interferes with YY1’s dimerization. FLAG-tagged YY1 and HA-tagged YY1 proteins were expressed in HeLa cells, the tagged YY1 proteins in nuclear extracts were co-incubated with anti-HA antibody for immunoprecipitation in the absence/presence of polyP_700_. The results showed that the FLAG-tagged and HA-tagged YY1 proteins interact as reported previously (Fig. 4C and 4D).^44^ However, this interaction is interrupted by polyP_700_ (Fig. 4C and 4D). In addition, we also triple-transfected the plasmids expressing FLAG-tagged YY1 and HA-tagged YY1 with an empty vector or PPK1 expressing vector in HeLa cells, nuclear extracts from these HeLa cells were immunoprecipitated with anti-HA antibody (Fig. S4A). The results showed that HA-tagged YY1 cannot dimerize with FLAG-tagged YY1 in nuclear extracts from PPK1-expressing cells (Fig. S4B), which is consistent with the findings observed upon polyP addition (Fig. 4D). YY1 is a known chromatin regulator controlling physical loop formation between enhancers and promoters in super-enhancer.^44^ Since YY1’s function to facilitate DNA interactions relies on its dimerization, we reasoned that polyP can also compromise DNA circularization facilitated by YY1. Indeed, purified YY1 has the ability to increase the rate of DNA circularization *in vitro* when a linear DNA template containing YY1 binding sites was incubated with purified YY1 followed by using a mobility shift assay (Fig. 5C and 5D). However, polyP_700_ blocks the circularization mediated by YY1 (Fig. 5C and 5D). These results indicate that polyP disrupts YY1’s function in mediating enhancer-promoter interaction.

**Figure 5.**
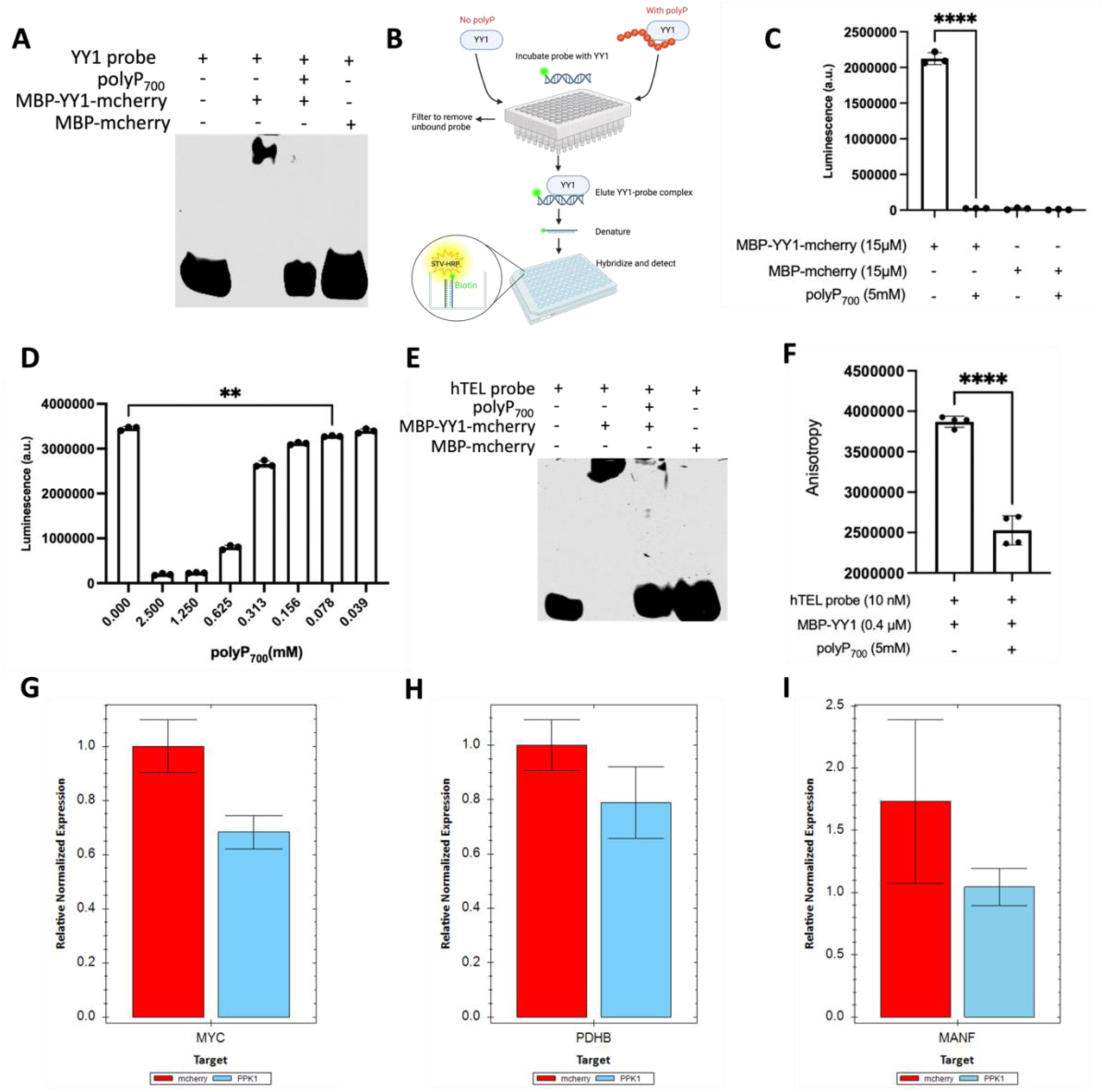
Polyphosphate inhibits YY1’s binding to DNA and affects gene regulation. (A) EMSA results using 4% native gel showing the binding of MBP-mCherry tagged YY1 to IR700Dye-labled YY1 binding motif after polyP_700_ treatment. (B) Schematic of the filter plate method used for assessing YY1 binding to its biotinylated DNA target. (C and D) Luminescent quantification of purified MBP-mCherry tagged YY1 binding to target DNA via the filter plate assay after treatment of 5 mM polyP_700_ (C) or the indicated concentration of polyP_700_ (D). Unpaired t test; ns, p > 0.05, ^∗∗^p ≤ 0.01, ^∗∗∗∗^p ≤ 0.0001, error bars ± standard deviation (n = 3). (E) EMSA results using 4% native gel showing the binding of MBP-mCherry tagged YY1 to IR700Dye-labled human telomere G4 structure after treatment of polyP_700_. (F) Fluorescence anisotropy for measuring the binding of MBP-YY1 protein with IR700Dye-labled human telomere G4 structures after polyP_700_ treatment. Unpaired t test; ns, p > 0.05, ^∗∗^p ≤ 0.01, ^∗∗∗∗^p ≤ 0.0001, error bars ± standard deviation (n = 3). (G, H and I) Quantification results of mRNA expression levels of *MYC* (G), *PDHB* (H) and *MANF* (I) genes in HeLa cells transfected with plasmids expressing either mCherry or mCherry-PPK1.

### Polyphosphate impairs YY1-DNA binding to interfere with its role in gene regulation

The specifical and efficient assembly of transcription apparatus at genomic sites is essential for gene expression regulation. YY1 specifically binds to a consensus motif, GCCGCCATTTTG,^41^ which is presented in both enhancer and promoter. This motif is important for YY1-mediated enhancer-promotor interactions and the formation of DNA loops.^44^ To further elucidate how polyP functions as a regulator to disrupt YY1’s role in gene regulation, we hypothesized that polyP blocks YY1’s ability to mediate DNA interaction by interfering not only with its dimer formation but also with its binding to the DNA consensus sequence. We tested this hypothesis using both traditional gel electrophoretic mobility shift assay (EMSA) and a more sensitive YY1 filter plate method where only DNA probes bound to YY1 are retained and subsequently quantified via luminescent detection (Fig. 5B). Results from both assays showed that polyP_700_ significantly impairs YY1-DNA binding (Fig. 5A and 5C). In human cells and tissues, the whole-cell levels of polyP range from 25 to 120 µM;^24^ in human platelets polyP can reach up to 130 mM in dense granules.^12^ To test whether the polyP levels in human cells are sufficient to affect DNA binding, we examined the effect of a gradient levels of polyP on DNA binding using filter plate. The results showed that polyP exerts stronger effect on DNA binding at higher concentrations, with as low as 78 µM of polyP still significantly impair DNA binding (Fig. 5D) This suggests that polyP may function as a regulatory molecule in various tissues and cells. More than 370,000 G4 structures, the four-stranded stable secondary structure of DNA, have been detected in human genome.^48^ These structures play important roles in various cellular processes in both normal and cancer cells, including replication and gene transcription.^45,48^ Importantly, YY1 dimers was recently found to directly bind to G4 structure to mediate long-range DNA looping and gene expression.^46^ We reasoned that polyP influences the interaction between YY1 and G4, and tested our hypothesis using EMSA and fluorescence anisotropy assay, as previously reported.^46^ Using a IR700Dye-labeled human telomere G4 (hTEL) probe, our results showed that polyP_700_ disrupts the strong binding of YY1 to the G4 structure (Fig. 5E and 5F). Furthermore, 1 mM polyP_700_ was sufficient to affect the binding, as indicated by EMSA (Fig. S4C). Results from fluorescence anisotropy showed that the impact of poly_700_ on YY1-G4 binding is significant (Fig. 5F). To explore whether polyP influences gene expression regulated by YY1, we assessed the messenger RNA expression levels of *MYC*, *PDHB* and *MANF* by using reverse transcription-quantitative PCR (RT–qPCR). All three genes were selected because Li et al.^46^ identified the presence of G4 structures and the occupancy of YY1 in their promoter regions. In addition, *MYC* was reported to be a tumor oncogene regulated by BRD4.^30^ The qPCR results demonstrated that the expression levels of *MYC*, *PDHB* and *MANF* were attenuated upon polyP production in HeLa cells (Fig. 5G, 5H and 5I). Taken together, these results indicate that polyP disrupts YY1’s binding to enhancer and promoter, thereby impairing its ability to mediate gene regulation.

### Short chain of polyphosphate impacts super-enhancer complexes

In mammals, polyP chains range from 2 to over 5000 orthophosphate units, depending on tissues and cell lines examined.^49,50^ Therefore, we investigated whether a short chain of polyP, polyP_100_, impacts the super-enhancer complexes. NuPAGE results showed that YY1, TAD-YY1, BRD4-PIR and MED1-PIR can also be modified by polyP_100_ (Fig. 6A, S3D, 6B and 6C), although the shifts of these proteins were smaller compared to those caused by polyP_700_. We investigated whether polyP_100_ affects the YY1’s recruitment of coactivators. The results showed that the droplets formed by YY1, BRD4-PIR and MED1-PIR are much smaller in the presence of polyP_100_ (Fig. S5), which consistent with effects of the long chain polyP (Fig. 4A and 4B). While the mCherry-YY1 still can colocalize with GFP-BRD4-PIR (Fig. S5A) or GFP-MED1-PIR (Fig. S5B). Next, we assessed whether polyP_100_ disrupts enhancer-promoter interaction mediated by YY1. First, we investigated whether polyP_100_ can also disrupt YY1 dimer formation. Indeed, with the addition of polyP_100_ in immunoprecipitation, the HA-tagged YY1 fails to form dimer with FLAG-tagged YY1 (Fig. 6D). In addition, we also investigated whether polyP_100_ inhibits the binding of YY1 to DNA consensus motif and G4 structure. The EMSA results showed that in the presence of polyP_100_, YY1 fails to bind to both DNA consensus sequence and hTEL G4 structure (Fig. 6E, 6G and S4D). Results from more sensitive assays, YY1 filter plate and fluorescence anisotropy, revealed that the impact of polyP_100_ on interactions between YY1 and DNA is significant (Fig. 6F and 6H). These results indicate that short chain of polyP can also work as a regulator in super-enhancer mediated gene expression.

**Figure 6.**
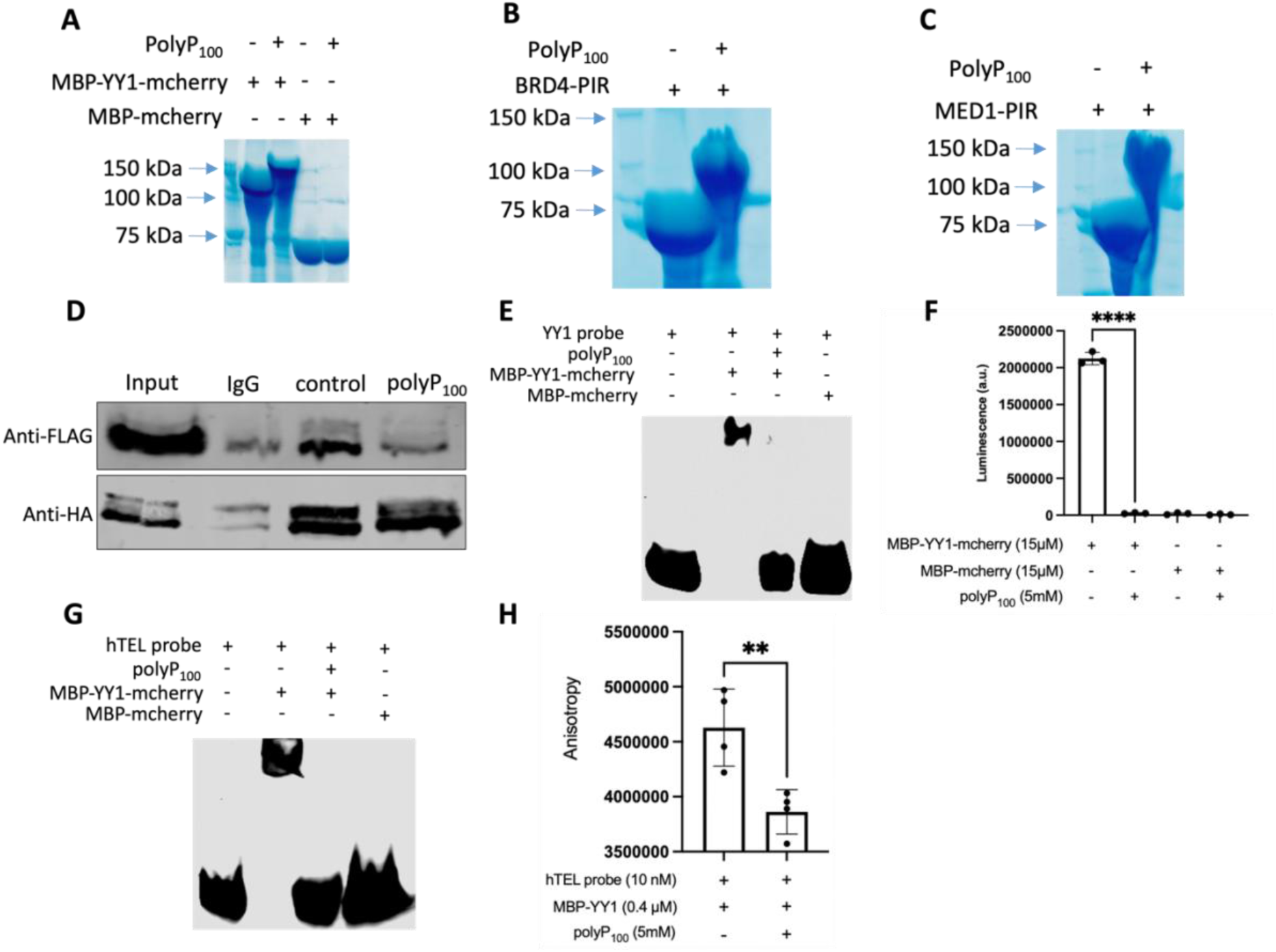
Short chain polyphosphate affects the functions of YY1, BRD4 and MED1. (A, B and C) Coomassie-stained NuPAGE analysis showing polyP_100_ modification on MBP-mCherry tagged YY1 (A), MBP-GFP tagged BRD4 PIR (B) and MED1 PIR (C). (D) Western blot analysis after SDS-PAGE showing the ability of polyP_100_ to impair co-immunoprecipitation of FLAG-tagged YY1 and HA-tagged YY1 proteins from nuclear lysates prepared from transfected cells using antibodies against FLAG or HA (n=3). (E) EMSA results using 4% native gel showing the binding of MBP-mCherry tagged YY1 to IR700Dye-labled YY1 binding motif after polyP_100_ treatment. (F) Luminescent quantification of purified MBP-mCherry tagged YY1 binding to target DNA via the filter plate treated with 5 mM polyP_100_. Unpaired t test; ns, p > 0.05, ^∗∗∗∗^p ≤ 0.0001, error bars ± standard deviation (n = 3). (G) EMSA results using 4% native gel showing the binding of MBP-mCherry tagged YY1 to IR700Dye-labled human telomere G4 structure with polyP_100_ treatment. (H) Fluorescence anisotropy for measuring the binding of MBP-YY1 protein with IR700Dye-labled human telomere G4 structures after treatment with polyP_100_. Unpaired t test; ns, p > 0.05, ^∗∗^p ≤ 0.01, error bars ± standard deviation (n = 3).

## Discussion

We describe here evidence that polyP interferes super-enhancer formation and therefore inhibits gene expression. PolyP acts as a versatile regulator of protein activity and cellular processes, influencing diverse biological functions. Super-enhancers, enriched with BRD4 and MED1, are vital for regulating gene expression across 86 human cell and tissue types.^29^ The conserved polyP interaction regions (PIRs) in BRD4, MED1 and YY1 are within intrinsically disordered regions (IDRs), underscoring the biological significance of polyP modifications. The PIRs feature His-rich, Lys-rich or PASK sequences and are targeted by polyP.^51^ IDRs of BRD4 and MED1 are essential for forming phase-separated condensates to compartment and concentrate the transcription apparatus at super-enhancer.^35^ The histidine-rich region of YY1, essential for droplet formation,^43^ is located near its coactivator-binding site, suggesting that polyP modification may obstruct coactivator recruitment or transcription apparatus compartmentation at super-enhancer. As a result of histidine and lysine polyP modification (HPM/KPM) of BRD4-PIR (Fig. 2C and 6B), MED1-PIR (Fig. 2C and 6C) and YY1-TAD (Fig. 3B and S3C), polyP directly interferes with co-localization of transcription factor YY1 and BRD4 (Fig. 4A) or MED1 (Fig. 4B) required for super-enhancer assembly. This disruption compromises the formation of super-enhancer complexes, impairing their functionality.

YY1 proteins form dimers to mediate the enhancer-promoter loop formation, contributing to super-enhancer-driven transcription.^44^ We have shown that polyP significantly disrupts YY1’s dimerization (Fig. 4D and 6D) and DNA looping (Fig. 4F). Our findings support the hypothesis that polyP acts as a potent regulator capable of modulating super-enhancers-driven transcriptional networks. This regulatory role is evidenced by polyP-mediated inhibition of nuclear localization (Fig. 3D-3G) and YY1-DNA binding (Fig. 5A-5F). Together, these effects collectively impair the expression of super-enhancer-regulated genes such as MYC, PDHB, and MANF (Fig. 5G-5I), highlighting polyP’s multifaceted impact on transcriptional regulation.

At higher salt concentrations, HPM/KPM of YY1, BRD4-PIR and MED1-PIR are disrupted (Fig. 2D and 3I), reducing its effect on phase separation (Fig. 2E and 3J). These indicate that the impact of polyP on proteins is salt-dependent, reflecting the non-covalent nature of HPM/KPM. This property aligns with prior findings and supports the reversable nature of polyP modifications.^27^ Notably, this feature has been exploited for purification of histidine-tagged proteins using immobilized polyP resin.^52^ Importantly, even at a concentration as low as 78 µM, polyP significantly disrupts protein-DNA binding, a value that aligns well with polyP concentration range reported in the literature,^24^ underscoring the physiological relevance of polyP as a regulatory molecule *in vivo*.

G4 structures are well-known promising anticancer targets.^48,53,54^ By disrupting YY1’s binding to G4 structures, polyP suppresses the expression of genes containing these structures (Fig. 5G-5I). This suggests that polyP may function as an anticancer regulator, mimicking the activity of G4-binding molecules, which are known to interfere with cancer-related transcriptional networks. In addition, disease-associated sequence variations are often enriched in super-enhancers.^29^ Cancer cells are heavily reliant on super-enhancers to drive dysregulated transcriptional programs critical for their identity and survival.^29,31–33^ By targeting super-enhancer complex components and the oncogenic transcription factor YY1 as evidenced in this study, polyP could reduce the expression of critical G4 containing oncogenes such as MYC (Fig. 5G). We anticipate that future studies will reveal additional roles of polyP as a modulator of dysregulation transcription in cancer. Conversely, long-chain polyP, known as a virulence factor in pathogens,^10,55^ may target host cell super-enhancers to impair immune responses during host-pathogen interactions. Thus, targeting bacterial polyP could serve as a promising therapeutic strategy against infectious diseases such as cystic fibrosis caused by *P. aeruginosa*.

BRD4 and MED1 are essential in forming super-enhancers by integrating transcription factors with the transcriptional apparatus, a process critical for gene regulation. However, BRD4 and MED1 protein levels decrease sharply in polyP overexpressing HeLa cell (Fig. 1D), while YY1’s localization to nucleus is only partially blocked (Fig. 3D and 3E). Importantly, we found that polyp targets four other transcription factors and additional four transcriptional activators.^21^ Therefore, we suspect that functions of these transcription factors are affected by polyP as well, particularly their roles in super-enhancers-driven transcriptional programs. As a result, this could reduce the need for or recruitment of BRD4 and MED1 to super-enhancer complexes. We believe that future studies will uncover more roles of polyP in modulating regulating gene expression, particularly with respect to other transcription factors and their involvement in transcriptional regulation. These discoveries could have significant implications for understanding polyP’s broader impact on cellular processes and disease mechanisms. In addition, our findings demonstrate that both long-chain (Fig. 2E, 4A and 4B) and short-chain (Fig. S5) polyP disrupt the phase separation of BRD4 and MED1. While polyP’s varying lengths in mammalian tissues and cells may enable polyP to exert differential effects on protein function, it is likely that polyP’s effect on super-enhancer is a general feature of mammalian gene control.

In summary, polyP emerges as a multifaceted regulator with profound implications for super-enhancer-driven transcription. By influencing enhancer-promoter communication and fine-tuning transcriptional bursts, polyP underscores its critical role in shaping the regulatory networks that govern gene expression.

## KEY RESOURCES TABLE

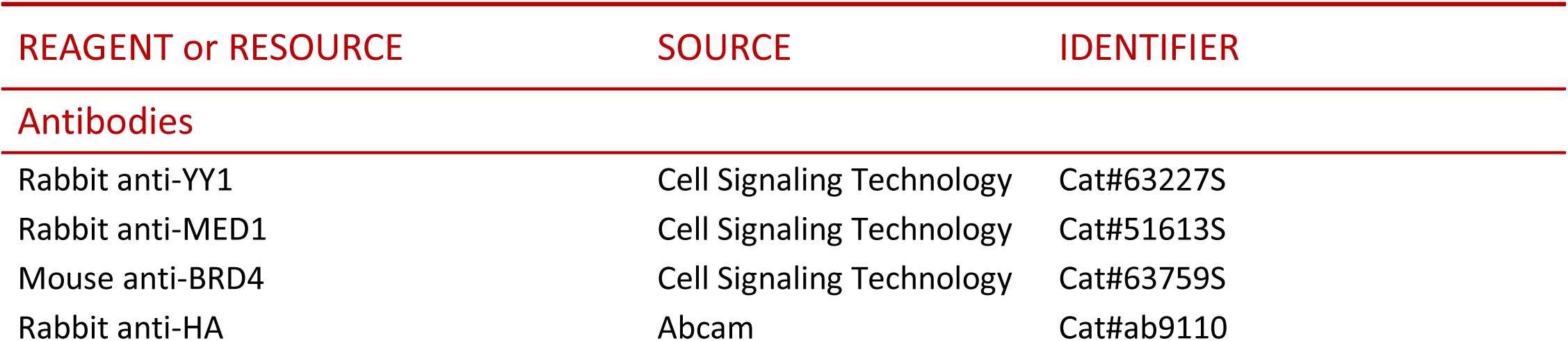
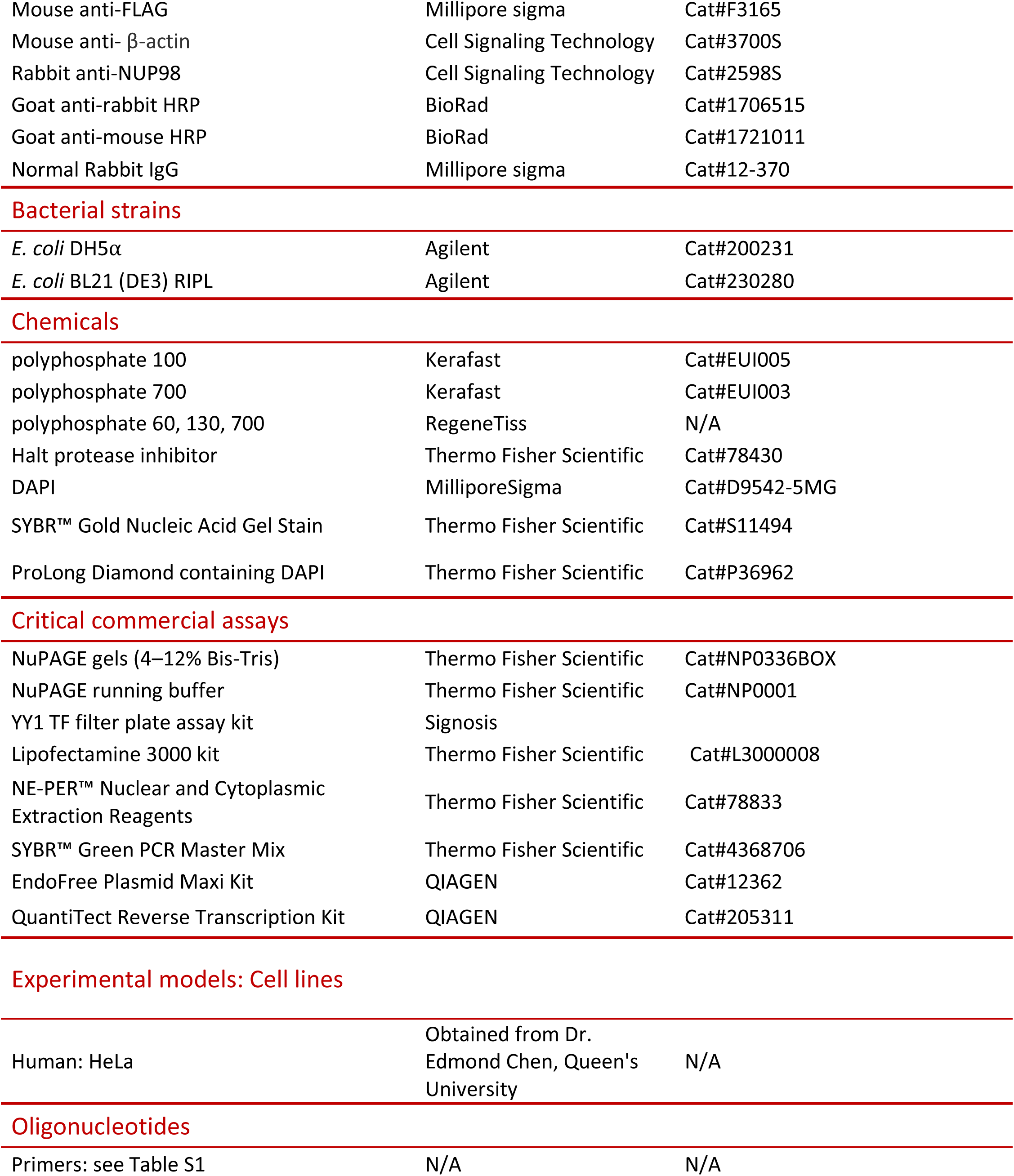
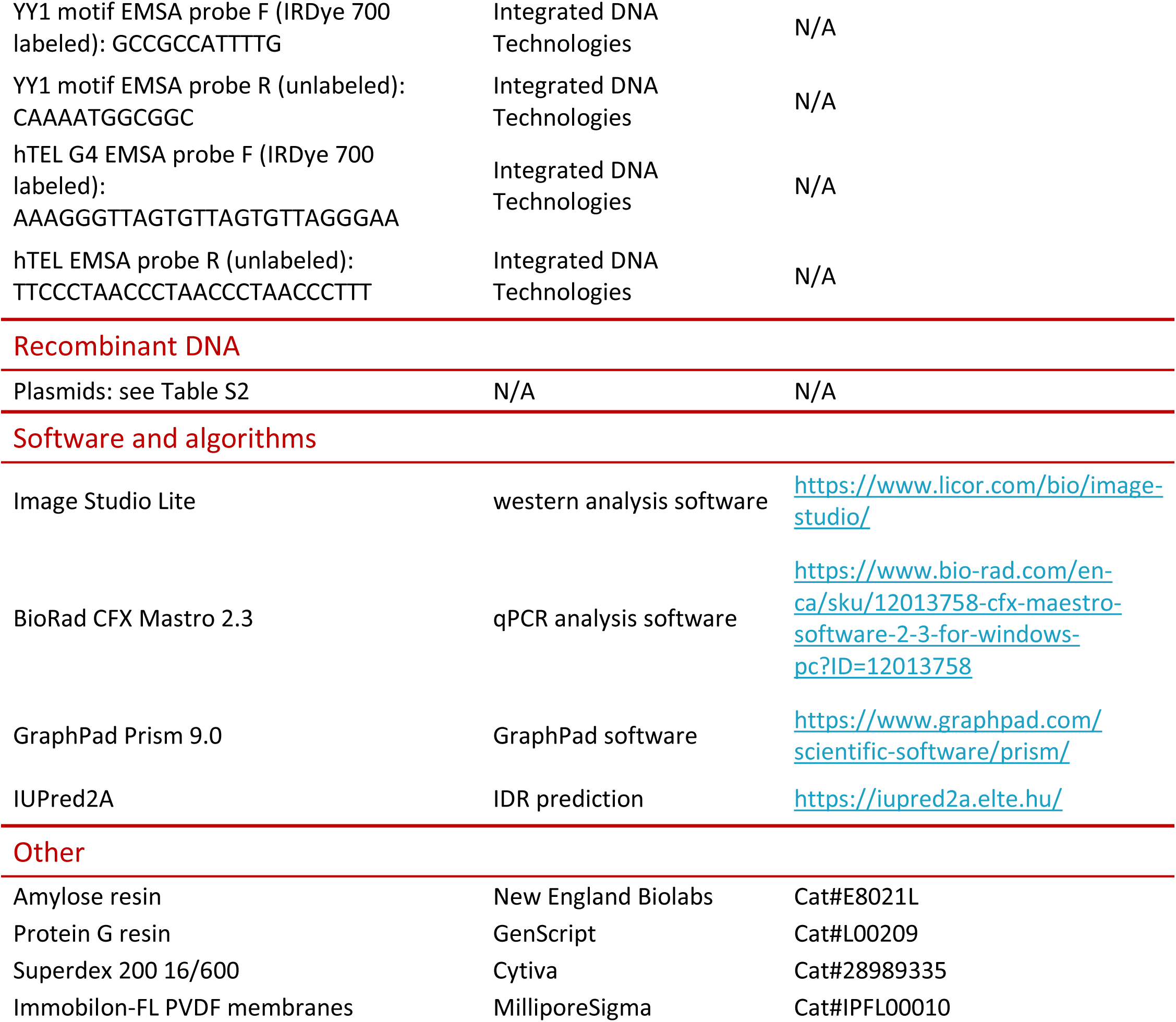

### Resource availability

#### Lead contact

Requests for resources and reagents and further information should be directed to and will be fulfilled by the lead contact, Zongchao Jia.

#### Material availability

Plasmids and reagents will be available upon request from the lead contact with a completed Materials Transfer Agreement.

### Experimental model and study participant details

#### Bacterial strains

Plasmid cloning was conducted in *E*. *coli* DH5α (Agilent, 200231). Recombinant protein expression was conducted in *E*. *coli* BL21(DE3) RIPL (Agilent, 230280). Bacteria were routinely grown in lysogeny broth (Bioshop, LBL407.5) supplemented with 50 μg mL^−1^ kanamycin or 100 μg mL^−1^ ampicillin depending on plasmids.

#### Cell lines

The HeLa cell line used in this study was obtained from the laboratory of Dr. Edmond Chen, Queen’s University. Cells were cultured at 37°C with 5% CO_2_ in Dulbecco’s Modified Eagle Medium (DMEM) with 10% fetal bovine serum. This cell line was routinely tested for mycoplasma contamination via the EZ-PCR Mycoplasma Test Kit (FroggaBio 20-700-20).

### Method details

#### Plasmids

A detailed description of the plasmids used in this study is available in Table S2. Oligonucleotide primers used are listed in Table S1.

#### HeLa transfection, protein extraction, and imaging

HeLa cells were seeded onto 6-well plates or glass coverslips pretreated with fibronectin in 24-well plates and transfected at ∼ 80% confluency using Lipofectamine 3000 (ThermoFisher L3000008) as per the manufacturer instructions. Cells were harvested at 48-hour post transfection.

To harvest protein extracts from 6-well plates, HeLa cells were washed three times with PBS, followed by pelleted and suspended in TNTE-FB lysis buffer (150 mM NaCl, 20 mM Tris pH7.5, 5 mM EDTA, 0.3% Triton X-100, 10 mM NaF, 40 mM β-glycerophosphate) supplemented with Halt protease inhibitor cocktail (ThermoFisher 78430). Then the cells were centrifuged for 3 min at 10,000 rpm to pellet debris. Supernatant was mixed 1:1 with 2x Laemmli loading buffer (120 mM Tris pH 6.8, 100 mM DTT, 20% glycerol, 4% SDS, 0.2% bromophenol blue) and boiled for 5 min prior to gel electrophoresis.

The nuclear and cytoplasmic extraction reagents (ThermoFisher 78833) were used to study the localization of YY1. Transfected HeLa cells were harvested with trypsin-EDTA, followed by cytoplasmic and nuclear proteins extracted as described by the assay instructions.

For imaging, transfected HeLa cells on coverslips were fixed with 4% paraformaldehyde in PBS for 10 min at room temperature, permeabilized with 0.1% Triton X-100, with three 5 min washed with PBS between each step. For polyP’s function on pre-formed YY1 droplets, permeabilized HeLa cells were incubated with 1 drop 50 mM polyP_700_. Coverslips were mounted with ProLong Diamond containing DAPI (ThermoFisher P36962). Protein localization or phase separation was observed via an Olympus IX83 microscope with 100x oil objective.

#### PolyP extraction and electrophoresis

HeLa cells transfected with mCherry or PPK1-mCherry were pelleted and lysed using GITC lysis buffer, and boiled at 95 °C for 10 min. PolyP was purified via silica spin column as described previously.^6^ The purified polyP eluate (15 µL) was then mixed with 15 µL of loading dye (10 mM Tris-HCl pH 7, 1 mM EDTA, 30% glycerol, bromophenol blue) and electrophoresed on 15.8% TBE-urea gels as described.^24^ The volume loaded for each sample was normalized with respect to the total protein content of the original cell lysate as determined via Bradford assay, in order to control for any variations in cell growth. PolyP was visualized via negative DAPI staining as reported previously.^56^ Briefly, gel was incubated in fixative solution (25% methanol, 5% glycerol, 50 mM Tris base, pH 10.5) containing 2 μg/mL of DAPI for 30 min at room temperature. Then, the gel was destained for 1 hour in fixative solution. Next, gel was exposed to 365 nm light on the UV transilluminator to induce photobleaching. This step required adjustable times (a few min) based on the amount of polyP. Finally, the gel was imaged using a GEL DOC 2000.

#### Protein expression, purification and size exclusion chromatography

Proteins were expressed using BL21(DE3) RIPL *E. coli* cells. Overnight cultures were inoculated into fresh LB at 1/100 dilution and grown at 37°C until OD_600_ = 0.6, at which point expression was induced with 1 mM IPTG, followed by growing the cultures at 16°C overnight. Cells were pelleted and resuspended in lysis buffer (50 mM Tris pH 8, 250 mM NaCl, 5% glycerol, 3 mM 2-mercaptoethanol, pH 8). Cells were lysed via sonication with a Branson instrument for 8 min, with cycles of 5s on and 15s off. Lysates were centrifugated at 18,000 rpm for 30 min. Clarified supernatant was applied to the amylose resin for MBP tags. After washing with lysis buffer for three times, proteins were eluted with 10 mM maltose.

The protein fractions were concentrated and further purified by FPLC using a Superdex 200 16/600 (Cytiva 28989335) size exclusion column. The column was equilibrated and run with no-salt buffer composed of 50 mM Tris, 50 mM MOPS, 1 mM EDTA, and 5 mM sodium bisulfite, final pH 7.5 to mimic NuPAGE gel running buffer, before the protein fractions were purified using the same buffer. After purified by FPLC, the proteins were concentrated and stored in 80°C.

#### *In vitro* polyphosphate modification assays

Polyphosphate modification assays was performed as described previously.^21^ Briefly, purified proteins or cell lysates were incubated with polyP for 30 min at room temperature. For salt effect experiments, the indicated concentration of NaCl was added to the buffer. Unless otherwise indicated, the polyP used throughout this paper was added to 5 mM final concentration. Samples were mixed 1:1 with 2x Lammli sample buffer and boiled at 95°C for 5 min. Samples were run on 4–12% Bis-Tris NuPAGE gels (ThermoFisher NP0336BOX) at 200 V for 45 min in a homemade 1X NuPAGE running buffer (50 mM Tris, 50 mM MOPS, 1 mM EDTA, 5 mM sodium bisulfite, and 0.1% SDS). The gel was used for Coomassie staining for western blot for further analysis.

#### Western blotting

Proteins were transferred to Immobilon-FL PVDF membranes (Millipore IPFL00010) for immunoblotting. The membrane was next blocked with 5% non-fat milk in TBST for 1 hour at room temperature with shaking. Membrane was then incubated with primary antibodies diluted in 5% non-fat milk or 5% BSA in TBST and incubated overnight at 4°C, with shaking. The primary antibodies used in this study are as follows: rabbit anti-YY1 (Cell Signaling 63227S), rabbit anti-MED1 (Cell Signaling 51613S), mouse anti-BRD4 (Cell Signaling 63759S), rabbit anti-HA (Abcam ab9110), mouse anti-FLAG (MilliporeSigma F3165), mouse anti-β-actin (Cell Signaling 3700S), rabbit anti-NUP98 (2598S), normal rabbit IgG (MilliporeSigma 12-370). In the next morning, the membrane was washed three times with TBST at room temperature for 5 min with shaking for each wash. Then incubated with secondary antibodies diluted in TBS at room temperature for 1 hour with shaking. The secondary antibodies used were goat anti-rabbit HRP (BioRad 1706515), goat anti-mouse HRP (BioRad 1721011). Followed by washing the membrane with three times with TBST at room temperature for 5 min with shaking for each wash. Blots were imaged using an LI-COR Odyssey CL-x.

#### *In vitro* phase separation

*In vitro* phase separation experiments of MBP-BRD4-PIR-GFP, MBP-MED1-PIR-GFP and MBP-YY1-mCherry were performed as described^21^ with slight modification. Briefly, the fusion proteins (approximately 4 mg/mL final concentration) and 5 mM polyP were added to a buffer containing 20 mM Tris-HCl pH 7.5, 1 mM DTT, and 10% PEG8000. After mixing, 5 μL of solution was immediately loaded onto a glass slide and covered with a coverslip, followed by imaged on an Olympus IX83 epifluorescence microscope.

#### EMSA

Proteins were prepared as above, and probes labeled with IRDye 700 was used to visualize binding. Labeled forward probes were based on the YY1 long consensus motif with sequence 5’-IRDye 700-GCCGCCATTTTG-3’, which was annealed to unlabelled reverse sequence 5’-CAAAATGGCGGC-3’ to create double-stranded probes; or based on human telomere G4 structure with sequence 5’ IRDye 700-AAAGGGTTAGTGTTAGTGTTAGGGAA-3’, which was annealed to unlabelled reverse sequence 5’-TTCCCTAACCCTAACCCTAACCCTTT-3’ in a buffer containing 10 mM Tris–HCl (pH 7.5), 100 mM KCl and 0.1 mM EDTA to form G4 structure. YY1 motif binding was conducted as described previously^21^ with slight modification. Briefly, 20 μL reactions contained 2 μL of 10X binding buffer (100 mMTris, 500 mM KCl, 10 mM DTT; pH 7.5), 2 μL of 25 mM DTT/2.5%Tween 20, 1 μL of probe (1 μM stock), 1 μL of 100 mM MgCl_2_, 10 μL protein (yielding approximately 10 μg per reaction), 1.5 μL of 100 mM polyP or water and 2.5 μL water. hTEL G4 structure binding was conducted as described.^46^ Briefly, 25 nM probe was incubated with 2 µM protein in a buffer containing 10 mM Tris–HCl (pH 7.5), 100 mM KCl, 10 μM ZnCl, 1 mM DTT, 3% glycerol and the indicated concentration of polyP. Reactions were incubated for 30 min at room temperature, then mixed with 4 μL of 6X orange loading dye. Samples were electrophoresed on 4% Native gel [for 40 mL mix: 6.67 mL 30% polyacrylamide stock (Polyacrylamide-BIS ratio = 29:1), 2 mL 1M Tris (pH 7.5), 7.6 mL 1 M Glycine, 160 µL 0.5 M EDTA, 200 µL 10% APS, 30 µL TEMED, 24.33 mL H_2_O] in TBE buffer [for YY1 motif binding: 5.5 g Bonic acid, 0.93 g EDTA and 0.4 g Tris in 1 L water, pH 7.0; for hTEL G4 structure binding: 40 mM Tris–HCl (pH 8.3), 45 mM boric acid and 1 mM EDTA]. Gels were run at 100 V for 30 min followed by imaged on an LI-COR Odyssey CL-x.

#### YY1 filter plate assay

The YY1 filter plate assay kit was purchased from Signosis (FA-0006), and assay was conducted as prescribed by Signosis instructions with 15 µM of purified proteins and indicated concentration of polyP in the reaction mix. The detection of bound probe was conducted using luminescence of an iD5 plate reader.

#### Fluorescence anisotropy

Fluorescence anisotropy measurements were conducted as previously described.^46^ Briefly, the binding assays were performed with 10 nM labeled hTEL G4 structure probe, 0.4 µM protein and 5 mM polyP or ddH_2_O in a binding buffer containing 10 mM Tris–HCl (pH 7.5), 100 mM KCl, 10 μM ZnCl and 1 mM DTT, followed by a 30-min incubation on ice. Fluorescence anisotropy was measured on an iD5 plate reader, with the excitation and emission wavelengths being 550 and 580 nm, respectively.

#### Co-immunoprecipitation

We received the pcDNA3_FLAG_YY1 and pcDNA3_HA_YY1 from Young’s Lab and the Co-immunoprecipitation was conducted as described^44^ with slight modification. Briefly, Nuclear extracts from HeLa cells single transfected with pcDNA3_FLAG_YY1 or pcDNA3_HA_YY1 and triple transfected with pcDNA3_FLAG_YY1, pcDNA3_HA_YY1 and mCherry or mCherry-PPK1 were extracted as described.^44^ To prepare beads for immunoprecipitation, 50 μL of protein G beads per immunoprecipitation was washed with 1 mL of blocking buffer (0.5% BSA in PBS) for three times, rotating for 5 minutes at 4°C for each wash. After centrifuging at 1000 g for 1 min, beads were resuspended in 250 μL of blocking buffer with addition of 5 µg of anti-HA or normal IgG antibodies and incubated for 1 hour at 4 °C with rotation. After incubation, beads were washed with 1 mL of blocking buffer for three times, rotating for 5 minutes at 4°C for each wash. Washed beads were collected via centrifuge before resuspended in nuclear extract.

For single transfection, equal amount of nuclear extracts from HeLa cells expressing pcDNA3_FLAG_YY1 or pcDNA3_HA_YY1 (100 µg each) were mixed with the washed beads with the addition of 5 mM polyP or ddH_2_O. For triple transfection, 100 µg of nuclear extracts from HeLa cells expressing mCherry or mCherry-PPK1 were added to washed beads. Beads were allowed to incubate with extract overnight at 4°C with rotation. The following morning, beads were washed with 1 mL of ice-cold wash buffer for five times, rotating for 5 minutes at 4°C for each wash. Washed beads were resuspended in 100 µL of elution buffer (0.1 M glycine, pH 2.5) and incubated at room temperature for 2 min before adding the 10 µL of neutralization buffer (1 M Tris-HCl, pH 8.5). Followed by boiled at 95 °C for 10 min. The Samples were used to test the immunoprecipitation results by western blot using anti-HA and anti-FLAG.

#### DNA circularization

DNA circularization was conducted as described^44^ with slight modification. Briefly, a PCR was run using pAW49 and pAW79 (received from Young’s Lab) as template to generate linear pieces of DNA. The PCR products were PCR purified (ThermoFisher K0702) and then digested with BamHI (NEB R3136) and PCR purified. The digested templates were used in the ligation assay. For ligation assay, approximately 2 nM DNA, 0.96 µg/L YY1, 5 mM polyP_700_ or ddH_2_O and 1 X T4 DNA ligase buffer (NEB B0202S) were incubated in a buffer composed of 10 mM Tris-HCl (pH 7.5), 100 mM KCl, 10 µM ZnCl_2_, 1 mM DTT and 10 mM NaCl at 20 °C for 30 min. Followed by adding 600 units of T4 DNA ligase (NEB M0202L) and incubating for 10 min at room temperature. After incubation, 30 mM EDTA, 1x NEB loading dye (NEB B7024S), 1 μg/μL of proteinase K (NEB P8107S) were added to the mix and heated at 65 °C for 5 min. Then samples were run on a 4%–20% TBE gradient gel for 3.5 hours at 120 V. The gel was stained with SYBR Gold (Life Technologies S11494) and imaged with a GEL DOC 2000.

#### RT-qPCR

Total RNA was extracted from HeLa cells transfected with mCherry or PPK1-mCherry using RNAzol^®^ RT (MilliporeSigma R4533) following the instruction. 1 µg of total RNA was used for reverse transcription assay using a reverse transcription kit (Qiagen 205311) for complementary DNA synthesis. RT-PCR was carried out using SYBR™ Green PCR Master Mix (ThermoFisher 4368706) on the CFX Opus 96 system (BioRad). Primers used for RT-PCR are listed in Supplementary Table 1.

#### Quantification and statistical method

Statistical details of experiments can be found in the figure legends. N represents biological replicates. Unpaired t test was performed in GraphPad Prism V9 to determine statistical significance. p values of less than 0.05 were considered significant.

## Acknowledgements

We thank Dr. Richard A. Young (Whitehead Institute for Biomedical Research and Department of Biology, Massachusetts Institute of Technology) for providing the pAW49, pAW79, FLAG-tagged YY1 and HA-tagged YY1 constructs; Dr. Susana De la Luna (Centre for Genomic Regulation, Barcelona) for providing the pEGFP YY1; Dr. Edmond Chan (Queen’s University) for providing mammalian cells. We thank Dr. Wenxi Feng for making a wood dark box for phase separation imaging and writing a python program for automatic phase separation imaging analysis. Our sincere thanks to Dr. Nolan Neville, whose earlier contributions on polyP were instrumental to this work. We thank Michelle Starkell and Andy Chen for assisting in western blotting and EMSA experiments. PolyP standards (P60, P130 and P700) were a generous gift from Dr. Toshikazu Shiba (RegeneTiss). Cartoon figures were created using BioRender. This work was supported by Natural Sciences and Engineering Research Council of Canada (RGPIN-2024-05181) and Cystic Fibrosis Canada (grant #1011927).

## Author contributions

Conceptualization, Z.J. and Z.Y.; methodology, Z.Y.; investigation, Z.Y.; writing – original draft, Z.Y.; writing – review & editing, Z.J. and Z.Y.; funding acquisition, Z.J.; supervision; Z.J.

## Supplemental information

Figure S1-S5

Table S1 and S2

