## Supplementary Information for "Polyphosphate as a Novel Regulator of Super-Enhancer Complexes: Disruption of Phase Separation and Gene Expression"

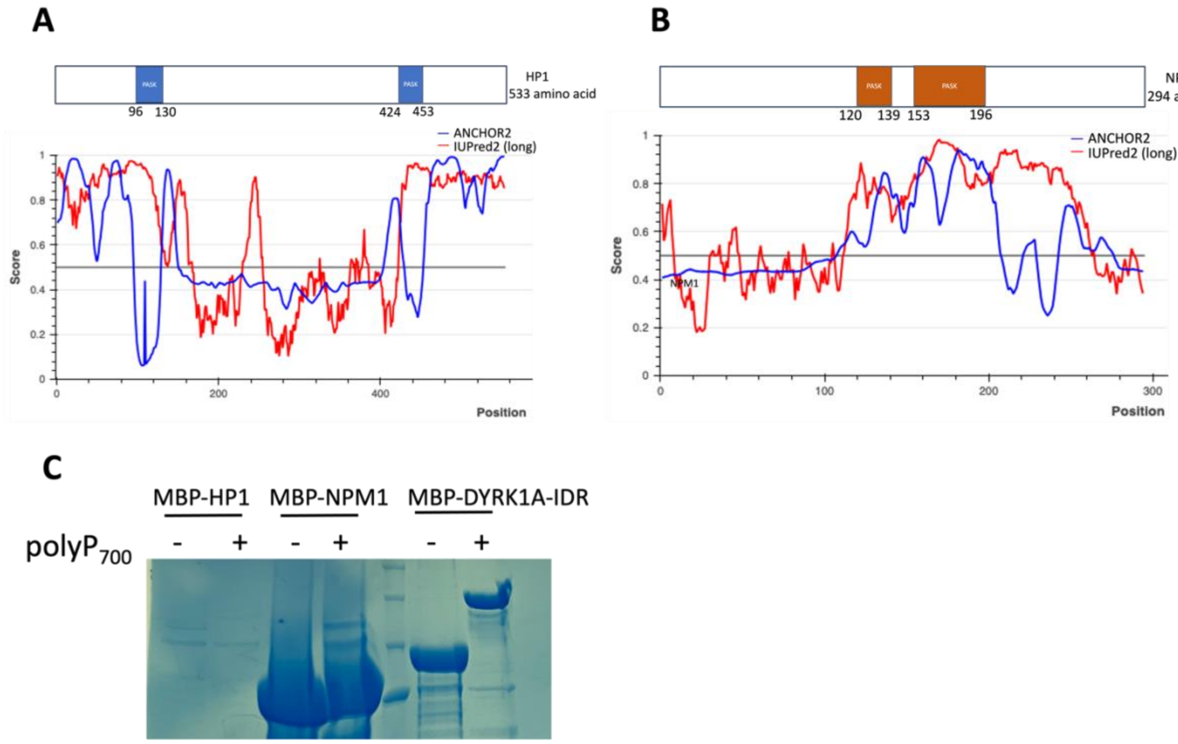


**Figure S1 polyphosphate cannot modify HP1 and NPM1**

(A and B) Domain graphs of HP1 (A) and NPM1 (B) PASK domains, IDRs prediction based on ANCHOR2 and IUPred2 algorithms. Scores >0.5 indicate disorder.

(C) Coomassie-stained NuPAGE analysis showing polyphosphate-mediated shift of purified MBP-tagged HP1 and NPM1 with $polyP_{700}$.

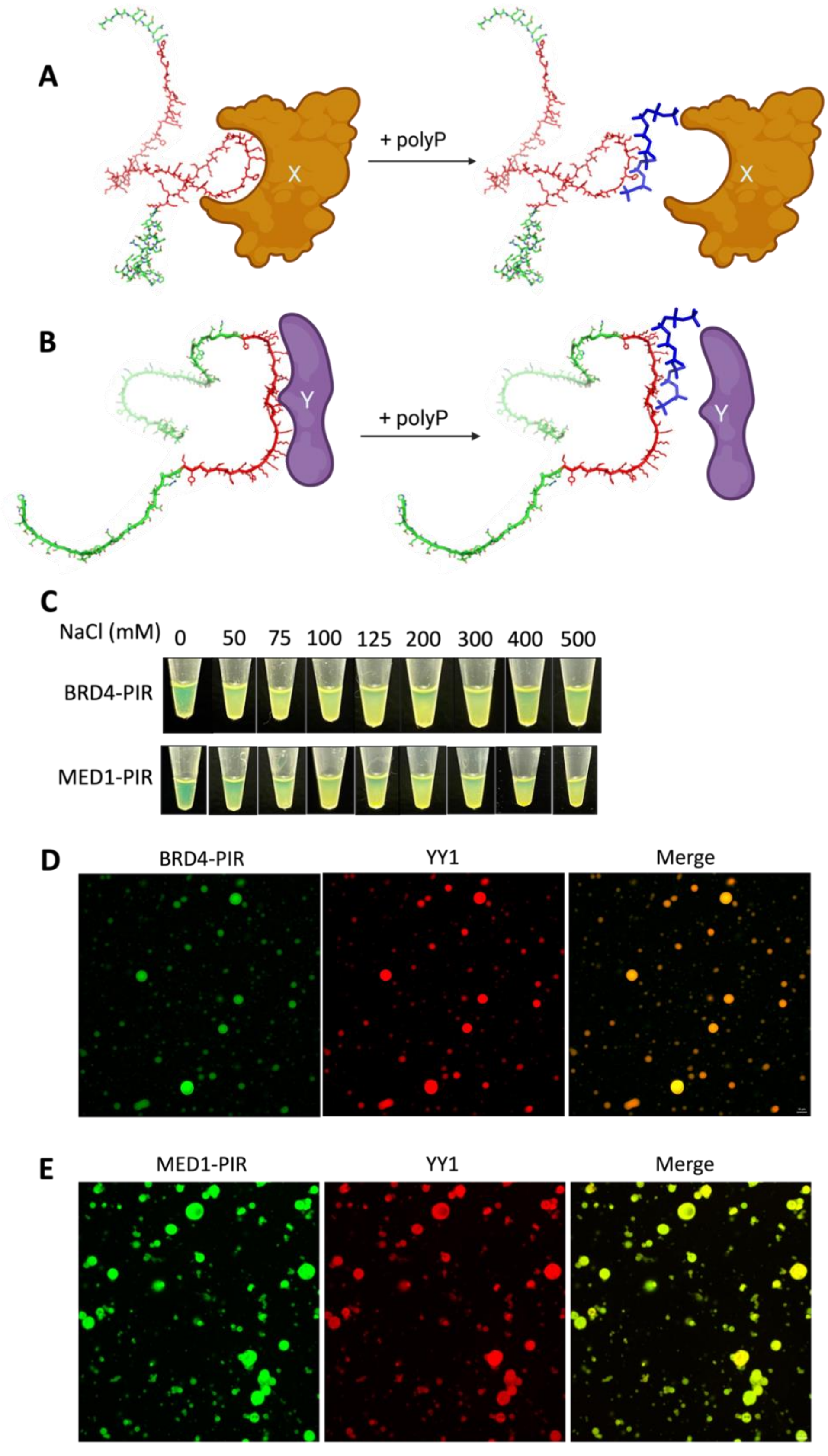
A
+ polyP
X
B
+ polyP
Y
C
NaCl (mM)
0
50
75
100
125
200
300
400
500
BRD4-PIR
MED1-PIR
D
BRD4-PIR
YY1
Merge
E
MED1-PIR
YY1
Merge

**Figure S2 Polyphosphate binds to the PIRs of BRD4 and MED1 and disrupts their liquid-liquid phase separation**

(A) The proposed model showing that polyP prevents the binding of BRD4 to its protein-binding partner (protein X) through PIR enriched in lysine. The PIR (amino acids 648-752) of BRD4 is highlighted in red.

(B) The proposed model showing that polyP prevents the binding of MED1 to its protein-binding partner (protein Y) through PIR enriched in lysine. The PIR (amino acids 1458-1581) of BRD4 is highlighted in red.

(C) Phase separation of purified MBP-GFP tagged BRD4-PIR and MED1-PIR with indicated concentration of salt. Tubes containing MBP-GFP tagged BRD4-PIR and MED1-PIR in the buffer containing PEG8000 (n=3).

(D and E) The ability of YY1 droplets to incorporate BRD4-PIR (B) or MED-PIR (C) proteins *in vitro*. The indicated MBP-GFP or MBP-mCherry fusion proteins were mixed in buffer containing PEG8000 and 100 mM NaCl. Indicated fluorescence channels are presented for each mixture (n=3).

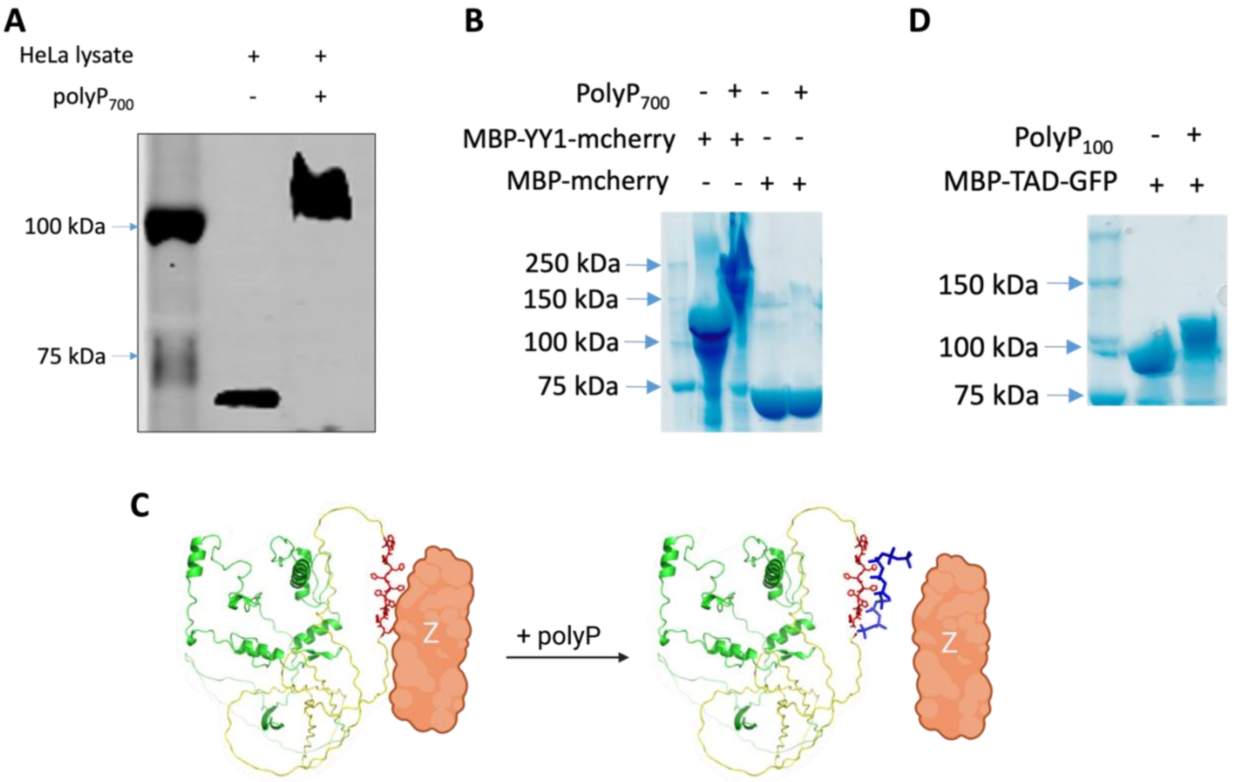

**Figure S3 Polyphosphate modifies TAD of YY1 through histidine-rich sequence**

(A) PolyP modification of YY1. HeLa cells lysate with/without the addition of $polyP_{700}$ were analyzed via NuPAGE followed by western blot with antibodies against YY1.

(B) Coomassie-stained NuPAGE analysis showing polyphosphorylation shift of purified MBP-mCherry tagged YY1 with $polyP_{700}$.

(C) The proposed model showing that polyP prevents the binding of YY1 to its protein-binding partner (protein Z) through PIR enriched in histidine. The histidine residues (amino acids 70-80) of YY1 are highlighted in red and YY1's transactivation domain (amino acids 1-154) is colored in yellow.

(D) Coomassie-stained NuPAGE analysis showing polyphosphorylation shift of purified MBP-GFP tagged TAD of YY1 with $polyP_{100}$.

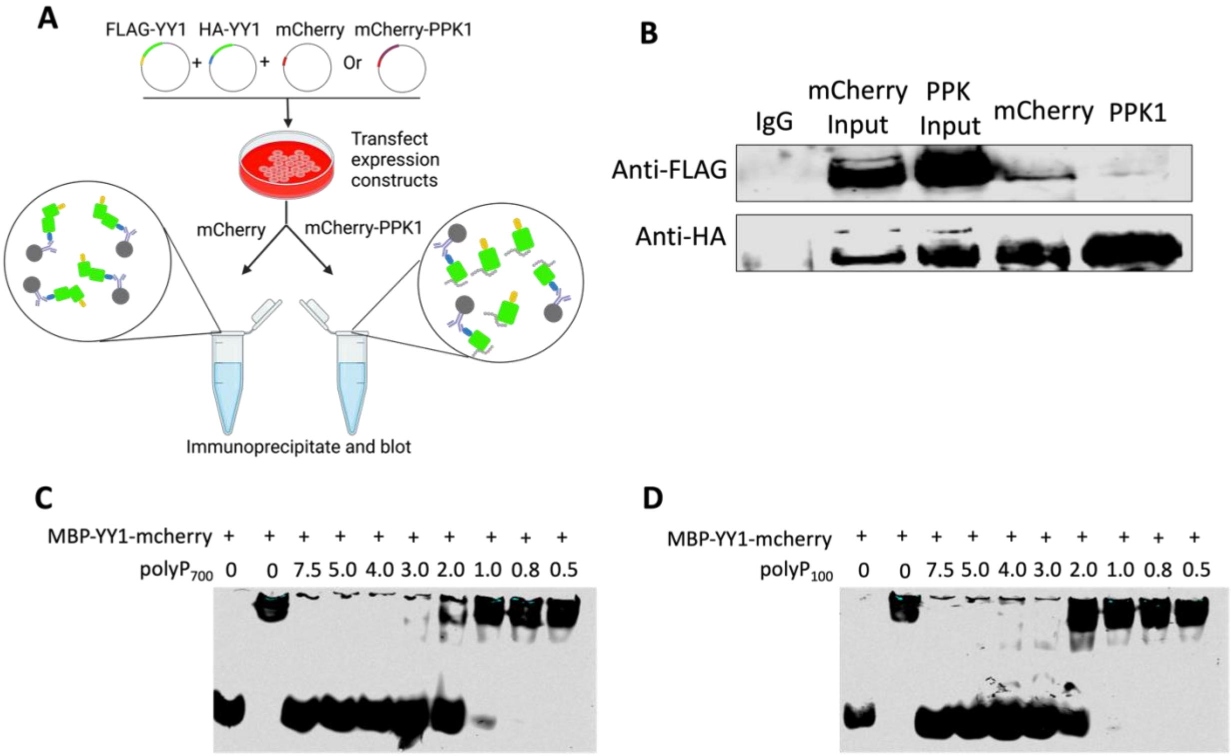


**Figure S4 Polyphosphate impacts on YY1 functions**

(A) A schematic depicting co-immunoprecipitation assay to detect YY1 dimerization in HeLa cells triple-transfected with FLAG-tagged YY1, HA-tagged YY1 and either mCherry or mCherry-PPK1 using HA antibody.

(B) Western blot analysis after SDS-PAGE showing the ability of polyP overproduction in HeLa cells to destroy co-immunoprecipitation of FLAG-tagged YY1 and HA-tagged YY1 proteins from nuclear lysates prepared from transfected cells using antibodies against FLAG or HA (n=3).

(C and D) EMSA results using 4% native gel showing the binding of MBP-mCherry tagged YY1 to IR700Dye-labled human telomere G4 structure after treatment with the indicated concentration of $polyP_{700}$ (C) and $polyP_{100}$ (D).

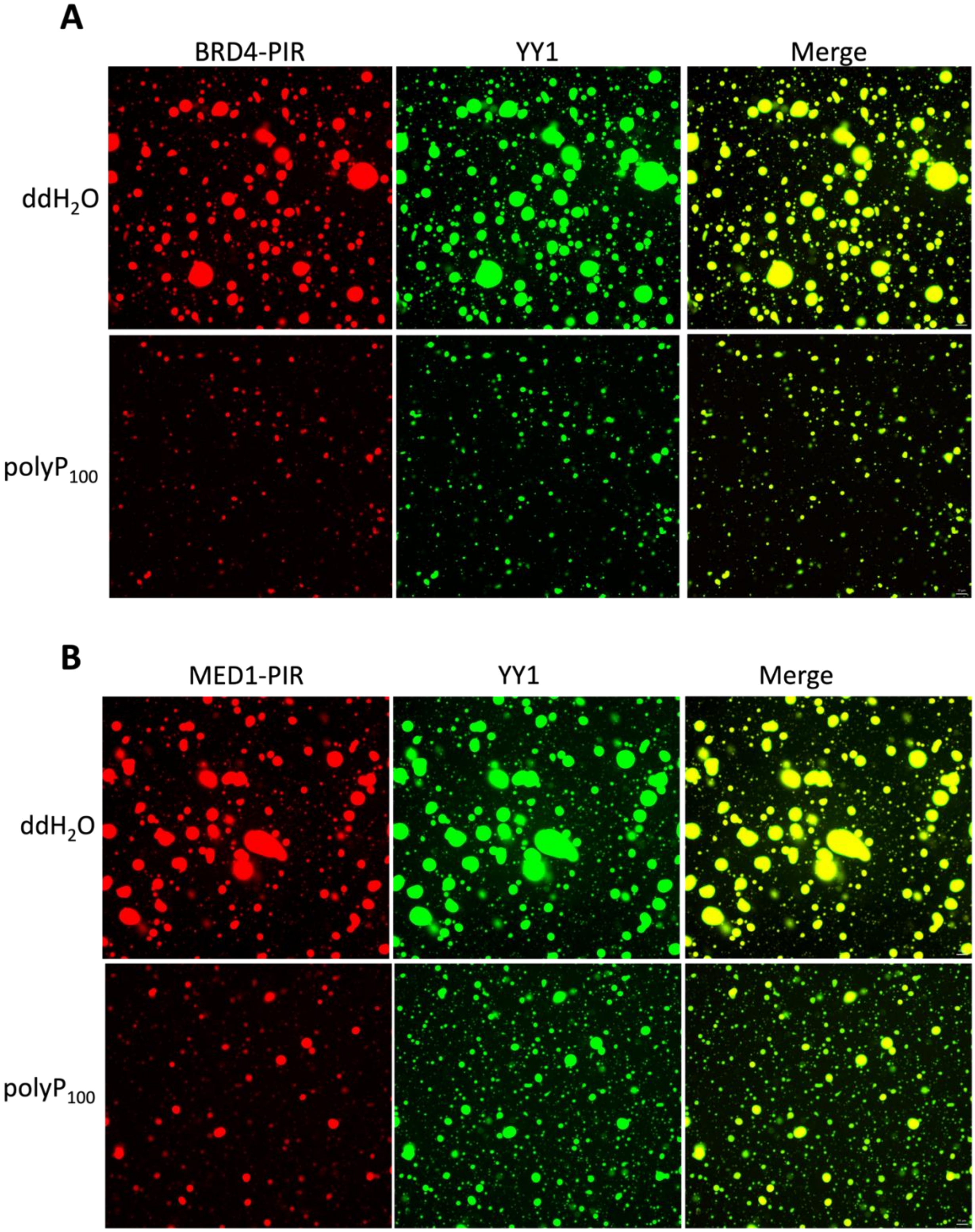


**Figure S5 Short chain of polyphosphate impact the phase separation of BRD4-PIR and MED1-PIR**

(A and B) $PolyP_{100}$ affects the droplet formation of BRD4-PIR (A), MED1-PIR (B) and YY1 (A and B); but not the ability of YY1 droplets to incorporate BRD4-PIR (A) or MED-PIR (B) *in vitro*. The indicated MBP-GFP or MBP-mCherry fusion proteins were mixed in buffer containing PEG8000 and 100 mM NaCl. Indicated fluorescence channels are presented for each mixture (n=3).

Table S1. Oligonucleotide primers used in this study. Relates to STAR Methods.

| Name | Sequence (5' to 3') |
|---|---|
| mCherry-PPK1-F | ATATAAGCTTCGCCACCATGAATACGCAGCAAGGACTTG |
| mCherry-PPK1-R | ATATGGTACCGCACGTGCGGTAAGCACCG |
| BRD4-PIR-GFP-F | ATATGGATCCATGAACCCCGACGAGATTGAAATCGACTTT |
| BRD4-PIR-GFP-R | ATATGAATTCCACAGGAGCCGGGGCCTGCTG |
| MED1-PIR-GFP-F | ATATGGATCCATGCAGAATCTGGACAGTGAAAGTGAGTCAGGC |
| MED1-PIR-GFP-R | ATATGAATTCATTCCCAATCAGGGCCACATCCATAAG |
| YY1-mcherry-F | ATATGGATCCATGGCGTCTGGTGACACCCTGTAC |
| YY1-mcherry-R | ATATCTCGAGCTGGTTGTTTTTCGCTTTCGCGTGG |
| YY1-TAD-GFP-F | ATATGGATCCATGGCGTCTGGTGACACCCTGTAC |
| YY1-TAD-GFP-R | ATATCTCGAGAGCAACGGTCACCAGGGTCTG |
| MANF-F | GGCGACTGCGAAGTTTGTAT |
| MANF-R | TTTGGTGGCTGCATCATCTG |
| PDHB-F | TCGAGGGCTGTGGAAGAAAT |
| PDHB-R | AATTCACAAATGGGCCGCAA |
| MYC-F | CTCCTACGTTGCGGTCACAC |
| MYC-R | CCGGGTCGCAGATGAAACTC |
| GAPDH-F | TTCGACAGTCAGCCGCATCTTCTT |
| GAPDH-R | CAGGCGCCCAATACGACCAAATC |

Table S2. Plasmids used in this study. Relates to STAR Methods

| Insert gene | MCS/Tag | Backbone vector | Description | Source |
|---|---|---|---|---|
| mCherry | N-MBP-TEV-insert-mCherry-C | pET16b | Control vector for expressing mCherry | Jia lab |
| GFP | N-MBP-TEV-insert-GFP-C | pET16b | Control vector for expressing GFP | Jia lab |
| YY1 | N-EGFP-insert-C | pEGFP-C1 | Human cDNA-derived *YY1* gene fusion to EGFP | Dr. Susana de la Luna |

| | | | | |
|---|---|---|---|---|
| PA PPK1 | N-insert-mCherry-C | pmCherry-N1 | *P. aeruginosa* PA14 *ppk1* gene fusion to mCherry | Jia lab |
| YY1 | N-MBP-TEV-insert-mCherry-C | pET16b | Human cDNA-derived *YY1* gene fusion to mCherry | This study |
| MED1-PIR | N-MBP-TEV-insert-GFP-C | pET16b | MED1 residues 1459-1581 from human cDNA-derived *MED1* gene fusion to GFP | This study |
| BRD4-PIR | N-MBP-TEV-insert-GFP-C | pET16b | BRD4 residues 648-752 from human cDNA-derived *BRD4* gene fusion to GFP | This study |
| YY1 | N-HA-insert-C | pcDNA3 | Human cDNA-derived *YY1* gene fusion to HA | Dr. Richard A. Young lab |
| YY1 | N-FLAG-insert-C | pcDNA3 | Human cDNA-derived *YY1* gene fusion to FLAG | Dr. Richard A. Young lab |
| YY1 binding sites | N/A | pAW49.pUC19 | N/A | Dr. Richard A. Young lab |
| Filler DNA instead of the YY1 motif | N/A | pAW79.pUC19 | N/A | Dr. Richard A. Young lab |
